# Creating an Ignorance-Base: Exploring Known Unknowns in the Scientific Literature

**DOI:** 10.1101/2022.12.08.519634

**Authors:** Mayla R. Boguslav, Nourah M. Salem, Elizabeth K. White, Katherine J. Sullivan, Michael Bada, Teri L. Hernandez, Sonia M. Leach, Lawrence E. Hunter

## Abstract

**Background:** Scientific discovery progresses by exploring new and uncharted territory. More specifically, it advances by a process of transforming unknown unknowns first into known unknowns, and then into knowns. Over the last few decades, researchers have developed many knowledge bases to capture and connect the knowns, which has enabled topic exploration and contextualization of experimental results. But recognizing the unknowns is also critical for finding the most pertinent questions and their answers. Prior work on known unknowns has sought to understand them, annotate them, and automate their identification. However, no knowledge-bases yet exist to capture these unknowns, and little work has focused on how scientists might use them to trace a given topic or experimental result in search of open questions and new avenues for exploration. We show here that a knowledge base of unknowns can be connected to ontologically grounded biomedical knowledge to accelerate research in the field of prenatal nutrition.

**Results:** We present the first ignorance-base, a knowledge-base created by combining classifiers to recognize ignorance statements (statements of missing or incomplete knowledge that imply a goal for knowledge) and biomedical concepts over the prenatal nutrition literature. This knowledge-base places biomedical concepts mentioned in the literature in context with the ignorance statements authors have made about them. Using our system, researchers interested in the topic of vitamin D and prenatal health were able to uncover three new avenues for exploration (immune system, respiratory system, and brain development), which were buried among the many standard enriched concepts, by searching for concepts enriched in ignorance statements. Additionally, we used the ignorance-base to enrich concepts connected to a gene list associated with vitamin D and spontaneous preterm birth and found an emerging topic of study (brain development) in an implied field (neuroscience). The researchers could look to the field of neuroscience for potential answers to the ignorance statements.

**Conclusion:** Our goal is to help students, researchers, funders, and publishers better understand the state of our collective scientific ignorance (known unknowns) in order to help accelerate research through the continued illumination of and focus on the known unknowns and their respective goals for scientific knowledge.

**Graphical Abstract:** 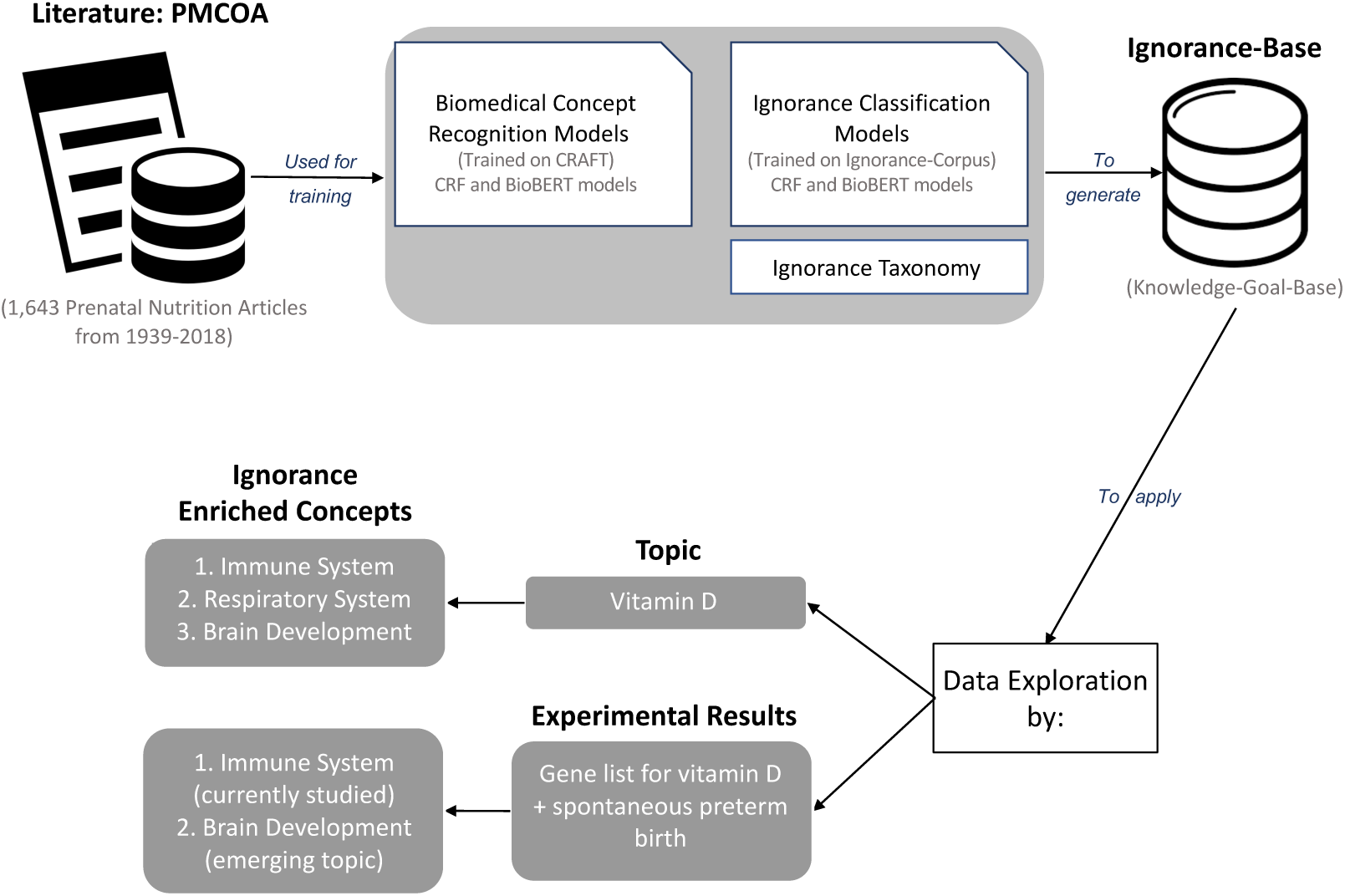

**Highlights:**

- We created the first ignorance-base (knowledge-base) to capture goals for scientific knowledge
- Our exploration methods provide analyses, summaries, and visualizations based on a query
- Ignorance enrichment provided fruitful avenues for future research
- Exploration by topic in vitamin D found three avenues to explore
- Exploration by experimental results for vitamin D and preterm birth found an emerging topic

## 1. Introduction

Research begins with a question. It progresses through accumulating knowledge such that a previously unexplored subject (an unknown unknown) becomes an active research area exploring the questions (known unknowns), until a body of established facts emerges (known knowns) [1, 2, 3]. We aim to help illuminate this process using biomedical natural language processing (BioNLP) to identify, categorize, classify, and explore known unknowns while highlighting their entailed goals for scientific knowledge (*i.e.*, actionable next steps). These known unknowns are discussed in the scientific literature as statements about knowledge that does not exist yet, including goals for desired knowledge, statements about uncertainties in the interpretation of results, discussions of controversies, and many others; collectively we call them **statements of ignorance**, borrowing the term from Firestein [1] and our prior work [4]. Our goal is to help researchers find the most pertinent questions to ask. For example, “these inconsistent observations point to the complicated role of vitamin D in the immune modulation and disease process” (PMC4889866) is a statement of ignorance. The entailed knowledge goal is to determine the correct role of vitamin D in the immune modulation and disease process by creating novel methods or conducting new experiments to study the complicated role. We also used **biomedical concept recognition**, the identification of biomedical vocabulary terms from ontologies or controlled vocabulary in text, to understand the biomedical subjects of these known unknowns. In the above example, these concepts include “vitamin D” and “immune”. Therefore, we aim to reveal these statements of ignorance, the entailed knowledge goals, and the entailed biomedical concepts to help students, researchers, funders, and publishers better understand the state of our collective scientific knowledge and **ignorance** (known unknowns).

While these ideas and methods are generally applicable across biomedical research, we chose to focus on the prenatal nutrition field. Due to ethical and legal considerations and complexities in studying pregnant mothers and fetuses, the field of prenatal nutrition is understudied and poised to benefit from the identification of questions that are well studied in other fields [5, 6, 7, 8]. Fetal development is a critical period and exposure to nutrition has a lifelong impact [9]. For example, the micronutrient vitamin D is very important for maternal and fetal health, affecting the immune and musculoskeletal systems, neurodevelopment, and hormones [10, 11, 12, 13, 14] (see Figure 1). Abnormal vitamin D levels can lead to gestational diabetes mellitus, preterm delivery, frequent miscarriages, adipogenesis, pre-eclampsia, obstructed labor, Cesarean sections, reduced weight at birth, respiratory issues, postpartum depression, and autism [10]. If we can identify the known unknowns or questions raised, even just with regard to the role of vitamin D, then we can search other fields for answers to inform the design of future studies. The prenatal nutrition field is a good case study for these ideas because it contains a diverse literature and a variety of studies from all over the world. Thus, applying an ignorance-based approach to this area is likely to generalize beyond prenatal nutrition, and more specifically to facilitate new interdisciplinary interactions that could advance the study of an underserved population and potentially help accelerate research to benefit mothers everywhere.

**Figure 1:**
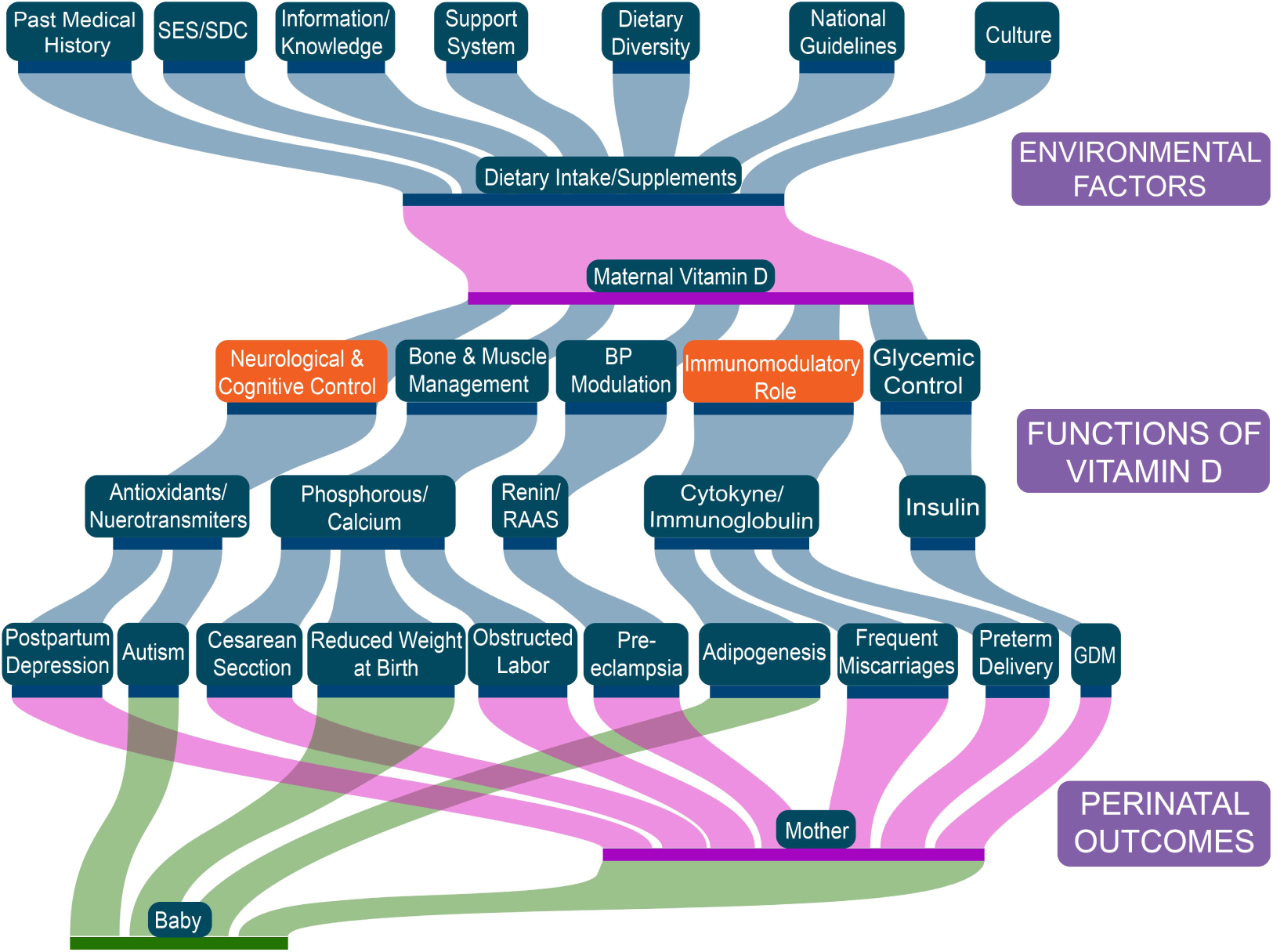
Relationship between society, maternal nutrition (vitamin D), and the effects on mother and offspring: a Sankey diagram created based on Figure 3 from [10]. The orange color represents the findings from the exploration methods that the concepts related to brain development and immune system are enriched in ignorance statements and possible novel avenues to explore. SES/SDC = socioeconomic status/sociodemographic characteristics; BP = blood pressure; GDM = gestational diabetes mellitus.

This work provides the necessary methods and tools to create a knowledge-base containing representations of known unknowns and their associated knowledge goals, an **ignorance-base**. This architecture allows scientists to explore the landscape of ignorance around a topic or a set of experimental results at scale and to find insights about related concepts across disciplines, resulting in an accelerated and interdisciplinary research process. (A list of the formal terms we have introduced here and their definitions are shown in Table 1.) We highlight its power by providing analyses, summaries, and visualizations that help researchers find knowledge goals to pursue in future work. Such an automated system could be useful to a wide variety of scientific stakeholders ranging from graduate students looking for thesis projects (*e.g.*, [15]) to funding agencies tracking emerging research areas (*e.g.*, [16]). It could help facilitate interdisciplinary interactions amongst researchers by finding questions from another field that bear on a topic or a set of experimental results (*e.g.*, [17]). It could also help track the evolution of research questions over time as a longitudinal analysis (*e.g.*, [18]). Furthermore, automatically identifying questions would allow us to query existing databases for relevant information (*e.g.*, [19]). Thus, there is a need for such an automated system to capture questions or known unknowns.

**Table 1:** Term definitions.

| Term | Definition |
| --- | --- |
| Ignorance | community/collective/scientific known unknowns |
| Knowledge-base | a database of known information |
| Ignorance-base | a knowledge-base, created from the literature, with additional annotations for the sentences that are ignorance statements |
| Statements of ignorance | statements of incomplete or missing knowledge categorized based on the entailed knowledge goal |
| Knowledge goal | the next actionable step based on the given unknown |
| Biomedical concept classification/recognition | automatically identifying and mapping biomedical entities to ontologies |
| Ontologies | controlled vocabularies with specified relationships |
| Open biomedical ontologies (OBOs) | an effort to create standardized ontologies for use across biological and medical domains |
| Lexical cue | words or phrases that signify a statement of ignorance |
| Taxonomy of ignorance | a categorization of ignorance statements based on the entailed knowledge goal |
| Exploration by topic | automatically find statements of ignorance related to a topic from the ignorance-base |
| Exploration by experimental results | contextualize experimental results in terms of statements of ignorance from the ignorance-base to understand what questions may bear on them |
| Ignorance enrichment | a method to identify biomedical concepts that are over-represented in a set of ignorance statements as compared to all sentences, and thus may be a new promising avenue to explore in relation to the input topic |
| Ignorance-category enrichment | a method to identify ignorance categories that are over-represented in a subset of ignorance statements as compared to all ignorance statements in order to illuminate the types of knowledge goals to pursue and to map out how they change over time |

There is only one similar system, the COVID-19 challenges and direction search engine (COVID-19 search engine) developed by Lahav *et al.*, [20]. They focused on creating a search engine to help researchers find two known unknown categories, scientific challenges and directions, and compared their work to a standard PubMed search. The search engine provides a relevant (high-confidence) table of challenge or direction sentences based on an input query of Medical Subject Headings (MeSH) terms. MeSH is a controlled vocabulary that is part of the Unified Medical Language System from NLM used for indexing, cataloguing, and searching for biomedical information and documents [21]). Their known unknown categories were motivated by the fact that “research focuses on *fine-grained* specific challenges, *e.g.*, difficulties in functional analysis of specific viral proteins, or shortcomings of a specific treatment regime for children. Each challenge, in turn, is associated with potential directions and hypotheses” [20]. However, their work stopped at identification of such statements and did not identify the knowledge goals associated with them; in contrast, our explicit representations of knowledge goals provide the users with guidelines for next research steps. In addition, we address a broader set of known unknowns and knowledge goals.

The other main goal of our work is to provide scientists with tools to explore the landscape of ignorance surrounding a topic or set of experimental results. The COVID-19 search engine [20] can support queries of multiple MeSH terms, but does not permit investigation of other types of inputs such as experimental results. As for the output, their search engine did not go beyond the identification of relevant sentences to provide analyses, summaries, or visualizations. This limits the ways users can explore the outputs to prioritize relevant areas of research. Lahav *et al.*, [20] posed as future work “to build more tools to explore and visualize challenges and directions across science.” Thus, their system could benefit from prior work focused on knowledge goals [4], the addition of other input types such as experimental results, and methods to explore and visualize known unknowns.

Our work extends the functionality described in [20] by means of the first ignorance-base. We compare our ignorance approach to the COVID-19 search engine and standard methods. Adding in prior work [4] that identifies statements about unknowns based on their entailed knowledge goals (**ignorance taxonomy**) provides actionable next steps for the users. For instance, we describe the specific challenges discussed above as *difficult tasks*, with the corresponding knowledge goals to create new tools or methods to overcome the difficulties and shortcomings. We create the ignorance-base based on this ignorance taxonomy by extending our prior work [4] to create more robust and high-quality classifiers of ignorance statements. Like many other knowledge modelers [22], we chose to ground our biomedical concepts in the open biomedical ontologies (OBOs) [23, 24] instead of MeSH for reproducibility, interoperability, and to avoid pitfalls in the modeling of knowledge [22]. Using these ontologies yielded state-of-the-art biomedical concept classifiers [25]. As a result of interweaving the ignorance-base with biomedical concept expansion, our work supports researchers in querying the literature for known unknowns, either by topic or with a list of experimental results, and then connecting this work to other knowledge-bases (*e.g.*, PheKnowLator [26, 27]) to find additional information relevant to the knowledge goals. Our system’s ability to perform concept enrichment and ignorance classification simultaneously extends its reach far beyond prior work [20], allowing it to trace out connections across different publications and knowledge-bases for a more comprehensive picture of what is known and unknown about a given subject.

The second goal of our work is to extend prior work [20] by adding in analyses, summaries, and visualizations of the outputs to help a researcher find knowledge goals to pursue. To do so, we explored the most frequent and enriched biomedical concepts in the ignorance statements returned by the input query in comparison to all sentences. This helped narrow researchers’ search for a topic in vitamin D to find ignorance statements ripe for exploration with the concepts “feeding behavior”, “immune system”, “brain development”, and “respiratory system” (see Figure 1). To help the researchers understand the general landscape of unknowns surrounding the topic of vitamin D and focus on the most interesting types, we summarized the ignorance categories around the enriched concepts and mapped out how they changed over time. We extended our prior work [4] by defining a total of 13 ignorance categories in the literature. For example, the researchers could choose a topic that is a complete unknown (*indication of unknown or novel research topic or assertion*) or a topic where there were alternate existing hypotheses to explore (*indication of alternative research options or controversy of research*). We demonstrate how this approach can help track the emergence of new research areas or produce a longitudinal analysis showing how research questions evolve over time. This is informative not only for scientists, but also for funding agencies and publishers [18, 28, 29, 30, 31, 32, 16, 33, 34, 35]. Once researchers have chosen a topic and want to evaluate experimental results, our goal is to help them contextualize those results in terms of statements of ignorance and understand what questions may bear on them, either within the same field as the topic or outside it. To do this, we conducted the same analyses as the input topic but also added canonical analyses based on the experimental results. Our motivating example was a gene list connecting vitamin D and spontaneous preterm birth (sPTB) from the literature [36]. If vitamin D plays a role in preventing sPTB, it would be relevant to all women of childbearing age. By comparing our ignorance approach to the standard approach for a gene list (functional enrichment analysis, gene list coverage, and the findings from the paper), we found ignorance enrichment of the concept “immune system”, a topic also identified by the original authors, as well as a novel putative relationship with the concept “brain development”, which implicates the field of neuroscience as a place to look for answers. We provide the ignorance statements and suggest questions for future exploration.

### Statement of significance

| Problem or Issue | What is Already Known | What this Paper Adds |
| --- | --- | --- |
| No knowledge-base focused on scientific knowledge goals from the literature exists to provide insights for contextualizing topics and experimental results in our collective scientific ignorance. | Knowledge-bases exist to find information and contextualize experimental results in what is known. Known unknowns are important and studied under different focuses. One search engine exists to help scientists discover challenges and directions. | This study aims to create an ignorance-base focused on knowledge goals to provide insights to researchers interested in exploring a topic or contextualizing experimental results in the known unknowns. We found emerging and currently studied avenues ripe for future work. |

## 2. Related Work

Many **knowledge-bases** (*e.g.*, the Reactome Pathway Knowledge-base [37]) exist to capture the known knowns from domain experts, the scientific literature, and other data sources such as experimental results [22]. These knowledge-bases have a variety of applications [22], including finding and interpreting information based on a single input topic, such as a concept, or a set of input topics that may be related, such as those from experimental results. In both cases the researchers want to find “relevant” information based on their query. For example, graduate students or researchers interested in learning about the field of prenatal nutrition might consult a database of dietary supplements [38]. Or researchers might perform a functional enrichment analysis to characterize a list of genes associated with vitamin D and preterm birth by finding relevant known biomedical concepts [36]. Many knowledge-base applications provide analyses to help researchers find and prioritize relevant information, which is our goal with the ignorance-base. Thus, we model the ignorance-base after knowledge-bases.

The aim of the ignorance-base is to help researchers find the most pertinent questions. Researchers gain these skills in graduate school, where the goal is to identify and provide at least some solutions for a question that is unanswered. There are many books [39, 40, 41, 42, 43] and articles [15, 44, 45, 46, 47] discussing how to choose the most pertinent question or topic, and yet only one automated system has been developed, the COVID-19 challenges and directions search engine [20]. As explained above, our goal is to extend their work to provide the users with knowledge goals and insights based on their input query just as in the knowledge-bases. One of the main differences between our work and the COVID-19 search engine [20] is in the categorization of known unknowns. Lahav *et al.*, [20] classified known unknowns into two categories, namely challenge and research direction; most of the similar prior work, including ours [4], introduced more fine-grained categories.

The original linguistics phenomenon that sparked all these areas of research was hedging. Hedged statements in linguistics can be true or false to some extent [48]. Recognizing that scientific research articles included hedges, hedging was then defined more specifically within these articles as “any linguistic means used to indicate either a) a lack of complete commitment to the truth value of an accompanying proposition, or b) a desire not to express that commitment categorically” [49]. Hedging highlighted a focus on truth and facts. To help specify the levels of truth, research turned to uncertainty, and the ways that a writer can communicate what they do not know to the readers. One of the first attempts to understand uncertainty theoretically was for decisionmakers, especially for law [50]. Scientific uncertainty was defined as the “different kinds of potential error associated with descriptive scientific information” [50]. This resulted in a taxonomy of six categories of descriptive uncertainty: conceptual, measurement, sampling, modeling, causal, and epistemic, each characterized by its own kinds of errors. In the bioscience field specifically, prior work sought to explore speculative language by presenting many examples of the phenomenon and determining that it was feasible for humans to annotate [51]. They focused on expressions of levels of belief including hypotheses, tentative conclusions, hedges, and speculations. Others have recast this phenomenon as factuality, alluding to a continuum that ranges from factual to counter-factual with degrees of uncertainty in between [52]. Still others [53] coined the term meta-knowledge to encompass different types of interpretive information including confidence levels, hypotheses, negation, and speculation. They determined five categories of meta-knowledge including manner, source, polarity, certainty level, and knowledge-type. All of these works focused on these phenomena in relation to the current known knowledge (*i.e.*, how certain, speculative, hedged, factual, or meta the knowledge is). More recently research has refocused these categories on goals for future knowledge, anticipating the next actionable step research should take in future work [4]. For example, the statement “there can be a relationship between smoking and lung cancer” is uncertain [54], and also an ignorance statement. The knowledge goal is to gather more evidence to support the claim (*indication of proposed or incompletely understood research topic or assertion*). Boguslav *et al.*, [4] identified 13 categories of ignorance and showed preliminary evidence for this categorization. One aim of our work is to build on this foundation to show the value of categorizing the knowledge goals of known unknowns for the ignorance-base.

Other prior work related to known unknowns includes efforts to capture them through understanding the phenomenon [48, 49, 55, 56, 57, 58, 59, 52, 51, 60, 61, 62, 63, 64, 65, 66, 67, 68, 69, 70, 71, 72, 73, 74, 75, 76, 77, 78, 79, 80, 81, 82, 83, 84, 85, 86, 87], creating taxonomies where a hierarchy of terms is linked by specified relationships [88, 62, 89, 52, 90, 91, 92, 50] and ontologies specifying relationships among controlled vocabularies [93, 94, 95, 96], annotating literature to create corpora [97, 98, 58, 56, 99, 100, 66, 101, 72, 82, 102, 103, 104], and automating identification of unknowns through classification tasks [105, 106, 52, 60, 61, 63, 64, 65, 67, 68, 69, 70, 71, 73, 76, 107, 77, 78, 79, 108, 81, 109, 83, 84, 110, 111, 85, 87, 59, 112, 113]. Some efforts have also sought to capture unknowns completely by creating theoretical frameworks, determining if the task is feasible for humans to perform, and automating it [4, 114, 62, 115, 116, 71, 75, 117, 80, 57, 86, 118, 51, 4, 20]. Only one work has created a formal search engine [20], and we create the first knowledge-base (ignorance-base) with added analyses, summaries, and visualizations that are made possible by a more fine-grained categorization of known unknowns.

Grounding our ignorance-base in the open biomedical ontologies (OBOs) [23, 24] also made our added analyses, summaries, and visualizations possible. Ontologies are vital to knowledge-based biomedical data science because they describe a knowledge representation in a way that preserves the defnitions of biomedical entities and the relations between them [22]. Additionally, ontologies license “the ontological commitments a knowledge representation makes (*i.e.*, what it can or cannot describe), which inferences are possible within it, and, sometimes, which of those inferences can be made efficiently.” [22] Within the biomedical domain, ontologies are “community consensus views of the entities involved in biology, medicine, and biomedical research, analogous to how nomenclature committees systematize naming conventions” [22]. Knowledge-bases grounded in and created from community-curated ontologies provide significant advantages for reproducibility in scientific research, for interoperability, and for avoiding pitfalls in the modeling of knowledge [22]. Knowledge-bases grounded in terminological resources, including UMLS (Unified Medical Languages System which includes MeSH - Medical Subject Headings), SNOMED CT (Systematized Nomenclature of Medicine, Clinical Terms), and the National Cancer Institute Thesaurus, lack some aspects of a computational ontology [22].

The COVID-19 search engine [20] used MeSH terms from UMLS, which lacks a common architecture and thus produces mappings that do not meld their terms together consistently into a single system [23]. UMLS [21] combines many vocabularies based only on the identification of synonymy relations between terms, resulting in potential loss of the intended meaning of concepts and distortion of the relationships between them during ontology mapping [119]. However, it can be easily applied to most currently existing databases [119]. Another effort to support biomedical data integration was the OBO foundry [23, 24], which sought to establish a set of principles for ontology development. These principles maintain the intended meaning of concepts, reduce the number and redundancy of ontologies, and require the cooperation and coordinated work of ontology developers [119]. Many OBOs, especially the Gene Ontology [120], are “specifically devoted to representing the biological knowledge underlying the reuse of data within new research contexts: in other words, it defines the ontology that researchers need to share to successfully draw new inferences from existing data sets” [121]. The goal of our work, using the OBOs, and that of Lahav *et al.*, [20], using the UMLS, is to find new insights from the literature on existing biomedical concepts. The OBOs contain many more terms/classes and asserted (nontaxonomic) relationships than MeSH (*e.g.*, the Gene Ontology [122, 120, 23]). The OBOs are generally semantically richer and allow for more semantic/logical entailments [122, 120, 23, 123]. Further, systems were created to help integrate the ontologies (*e.g.*, BioPortal [124]). The downside of the OBOs is that they only integrate well with other databases derived from OBO Foundry ontologies [119]. We chose to use the OBOs because their richness and interoperability provide assurance that future work based on the OBOs can continue to build on our work. In time, the OBO model is also likely to remedy some of the flaws in the UMLS [119], allowing future work to combine these efforts at standardization.

Our main novel contribution is providing analyses, summaries, and visualizations to help researchers find the next areas to study (biomedical concepts) and pertinent questions to ask (ignorance statements). Note that our analyses are made possible by both the knowledge goal categorization of known unknowns [4] and the OBOs [23]. Lahav *et al.*, [20] posed future work to “build more tools to explore and visualize challenges and directions across science” [20]. This work is the first step towards those goals. Further, many knowledge-base applications for experimental results are used to provide the researchers with a “list of ‘interesting’ biomolecules” [125]. Functional enrichment analysis is the standard method for obtaining such lists and has become one of the most frequently used tools in computational biology [125, 126, 127, 128, 129]. For example, functional enrichment analysis provides valuable insight into a collective biological function underlying a list of genes “by systematically mapping genes and proteins to their associated biological annotations … and then comparing the distribution of the terms within a gene set of interest with the background distribution of these terms” to identify statistically over- or under-represented terms within the list of interest [125]. The set of enriched terms then describe some important biological process or behavior [125]. Our work aims to provide a similar list of enriched terms with regards to ignorance (ignorance enrichment and ignorance-category enrichment) based on an input topic or set of experimental results, and use it to create summaries and visualizations to help researchers narrow in on the next areas to study and the pertinent questions to ask.

The goal of our system is to use the ignorance-base and exploration methods to go beyond the usual reach of a search engine, namely to provide summaries and visualizations for the numerous sentences and articles returned from an input topic or set of experimental results. We used ideas and techniques from the field of document visualization, which “transforms textual information such as words, sentences, documents, and their relationships into a visual form, enabling users … to lessen their mental workload when faced with a substantial quantity of available textual documents” [130]. Gan *et al.*, [130] provided an overview of the field with design principles and examples. They discussed visualization techniques for both single document and document collection visualizations as well as vocabulary-based visualizations to visualizations of document similarity [130]. We utilize Tag Clouds [130] to visualize the frequency of both words and biomedical concepts to compare them for ignorance statements versus all sentences. Other vocabulary-based visualizations include Wordle, TextArc, and DocuBurst [130]. Visualizations based on semantic structure include Semantic Graphs and visualizations based on document content include WordTree and Arc Diagram [130]. Visualizations for collections of documents can illustrate document themes, document core content, changes over different versions, document relationships, and document similarity [130]. All of these can help researchers gain an overview of the entire collection [130]. This work provides preliminary visualizations to help researchers digest the output of the ignorance-base. Future work can add and evaluate more visualizations.

The ultimate goal of this work is to provide analyses, summaries, and visualizations of ignorance statements resulting from an input topic or set of experimental results. For an input topic, similar works are search engines including PubMed and the COVID-19 search engine [20]. We compare our results to them. For experimental results, methods for standard functional enrichment analyses (contextualizing experimental results) use knowledge-bases and ontologies [125, 126, 127, 128, 129], and natural language processing (NLP) tools over the biomedical literature [17, 131, 132, 133, 134, 135, 136, 137, 138, 139, 140]. Some of this prior work not only aimed to characterize genes but also to help define new research areas (*e.g.*, [17] as one of a few goals), generate new hypotheses (*e.g.*, [141]), find information about genes of unknown function and fill gaps in knowledge (*e.g.*, a preprint [139] using manual curation). Thinking beyond a gene list, if we consider pathway models as experimental results, tools exist to associate pathway models to the literature (*e.g.*, [142]) and some of these take uncertainty into account (*e.g.*, [60, 143]). These works however focus on confidence and relevance to current knowledge, respectively, rather than focusing on the role they play in future knowledge and explicitly representing statements of known unknowns. Thus, instead we compare our results to the standard functional enrichment analysis [144, 145] to highlight the difference and power of ignorance enrichment. We build upon all of this previous work to create an ignorance-base grounded in knowledge goals and OBOs to explore by topic and experimental results, providing researchers with tools and visualizations to explore the landscape of our collective scientific ignorance.

## 3. Methods

We created an ignorance-base grounded in knowledge-goals (ignorance) and OBOs to provide analyses, visualizations, and summarizations to researchers to help them find pertinent questions to explore in future work. We combined the best-performing ignorance classifiers (extending the work of [4] to create a corpus of 91 articles) with state-of-the-art biomedical concept classifiers [25] to create the ignorance-base and explore it by a topic and by experimental results. The ignorance-base can be queried by ontology concepts, ignorance categories, specific lexical cues, or any combination of the three. We compared our results to standard methods and to the COVID-19 search engine [20].

The rest of this section is organized into the following subsections:

1. Creating the ignorance-base
2. Exploration by topic
3. Exploration by experimental results

### 3.1. Materials

The inputs for all systems were scientific prenatal nutrition articles. We used full-text articles from the PubMed Central Open Access (PMCOA) subset of PubMed [146], allowing us more data beyond the abstract and the ability to share it publicly. 1,643 prenatal nutrition articles (1939-2018) were gathered from querying PMCOA for 54 regular expressions (keywords such as {prenatal, perinatal and antenatal} paired with keywords like {nutrition, vitamin and supplement} determined in consultation with a prenatal nutrition expert, Teri L. Hernandez. All articles were provided in XML format, which was parsed and converted to text format using a script in Java. All subsequent computation was implemented in Python 3, with its associated packages. The continued annotation effort used Knowtator [147] and Protege [148] as in previous work [4], allowing the ignorance taxonomy (see Table 2) to be easily browsable like an ontology, and helping the annotators select the correct level of specificity for each lexical cue. The classification frameworks and models were also from our previous work [25, 4].

**Table 2:** Ignorance Taxonomy: definitions, knowledge goals, example cues, and total cue count. The categories in bold are only narrow categories. Abbreviations are in italics.

| **Ignorance Category** | **Definition** | **Knowledge Goal** | **Example Cues** | **Total Cues** |
| --- | --- | --- | --- | --- |
| **indication of *answered* research *question*** | A statement of a goal or objective of a study that is attempted or completed during the study. | to find the answer(s) in the article; determine if the question(s) is (are) fully answered in the article | aim, goal, objective, our study, sought, to determine | 64 |
| indication of *unknown* or *novel* research topic or assertion | A statement that indicates something is not known (a lack of information), or information is presented for the first time (new or novel) and a significant amount of research is needed; not a statement about the absence of something. | to explore the unknown further to gain any insights | could not find, don't know, elusive, not...established, uncertain, still unclear | 155 |
| indication of *explicit* research *inquiry* | An explicit statement of inquiry (with a question mark or question word such as how, where, what, why). | to find answers to the question and/or discover methodologies that will help answer the question | ?, what, where, wondered, why | 19 |
| indication of proposed or *incompletely understood* research topic or assertion | A positive or negative statement proposing a possible/feasible explanation for a phenomenon on the basis of limited evidence as a starting point for further investigation OR a statement that information is needed to support an assertion or claim, including both positive and negative statements. Either a statement that some evidence already exists, explaining how current findings support previous work, adding confidence to a claim OR a statement that information is limited, more research is needed or is ongoing including limitations – biases or shortcomings related to the study design and execution. | to gather more evidence to support the claim OR conduct more research to determine the validity of the claim; complete the partial picture; consider the shortcomings and biases for the next experiment and how it can be addressed. | a good understanding, believe, evidence...limited, has been suggested, hypothesis, no studies, possibly, preliminary stage, remains under investigation, still being discovered, support, trend | 797 |
| indication of *indefinite relationship* among research variables | A statement about a connection, link, or association between at least two variables; connectedness between entities and/or interactions representing their relatedness or influence. | to confirm the connection, link, or association between variables; determine the full underlying relationship between variables | affect, associated, correlate, factor, influence, interact, link, pattern, tend | 198 |
| indication of *largely understood* research topic or assertion | A statement staking a claim to the most likely explanation, relationship, or phenomenon; assumes that there is a good chance this understanding is correct. | to determine if the most likely option is correct or if another option is more feasible | almost all, assumed, concluding, evident, it is clear, most likely, thus | 202 |
| **indication of *anomalous* or *curious* research finding** | A statement of a surprising result, conclusion, observation or situation; the researchers were not expecting the result, conclusion, observation or situation but are intrigued by it. | to explore the surprising result, conclusion, or situation more and determine if the result, conclusion, observation, or situation is repeatable | appeared to be, interestingly, noteworthy, surprisingly | 113 |
| indication of *alternative* research options or *controversy* of research | Either an explicit statement of multiple (at least two) choices, actions, approaches, or methods that need to be experimentally determined, including statements with an implied second option, such as “whether”. This includes a statement of disagreement amongst researchers OR a lack of consensus OR at least two possible answers presented as results from different researchers, usually in reference to previous results and stated when results contradict or disagree with each other. | to determine the correct option or a better option and if there are disagreements, to determine the truth to break any disagreements | cannot rule out, claims, has been challenged, whether, whilst | 221 |
| indication of *difficult* research *task* | A statement of something not easily done, accomplished, comprehended, or solved; or a complicated thing with a multitude of underlying pieces or parts; heterogeneity; excludes medical complications. | to create methods to study the complicated system and to better understand any piece of the complicated system; potentially requires new experiments or better techniques | not feasible, remains...challenge, variability, rarely able to | 98 |
| indication of research *problem* or *complication* | A statement of issues, problems, mistakes, or medical complications that are cause for anxiety and/or worry. | to determine the gravity of the concern and determine if it needs to be dealt with before the next experiment or study | issue, error, insufficient, lack of reproducibility, publication bias, underestimated | 98 |
| indication of *future* research *work* | A statement of extensions, including next steps, directions, opportunities, approaches, or considerations of the described work that may be implemented at some future time point. This also includes a statement of suggestion or a proposal as to the next best course of action, especially one put forward by an authoritative body; advice telling someone the best action to take. | to determine the next course of action based on this future work proposal | additional research, are needed, continue to explore, further study, more...studies, recommend, warrants, worthy of closer attention | 258 |
| indication of *future* research *prediction* | A statement of extrapolation of given data into the future and/or from past observations, without reference to next steps. | to run the simulation or experiment to determine if the prediction is correct; publicize the outcomes of the study to the correct people | allow, expect, if so, serve as a basis, will | 27 |
| indication of *important consideration* for future research work | A statement calling for attention including an action needed to be taken immediately or information that needs to be disseminated immediately OR is critical: being in or verging on a state of crisis or emergency OR urgently needed OR absolutely necessary. | to take the urgent action ASAP or distribute the knowledge ASAP | call for action, cautious, crucial, emphasis, global problem, high on the agenda, necessary, relevant to note, vital | 263 |

To connect the ignorance statements to the biomedical concepts, the ignorance-base was built upon the PheKnowLator knowledge graph (Phe-KnowLator v3.0.2 full subclass relationsOnly OWLNETS SUBCLASS purified NetworkxMultic DiGraph.gpickle), which semantically integrates eleven OBOs [26, 27]. For exploration by topic, we compared our results to a PubMed literature search and the COVID-19 search engine [20]. For exploration by experimental results, the gene list (our motivating example of experimental results) was gathered from a PMCOA article (PMC6988958) [36]. We also used DAVID (a tool for functional annotation and enrichment analyses of gene lists) [145] as a standard approach for functional enrichment analysis to compare to ignorance enrichment.

Computation used a contemporary laptop (MacBook Pro) and an NIH-funded shared supercomputing resource [149] that included:

- 55 standard compute nodes with 64 hyperthreaded cores and 512GB of RAM
- 3 high-memory compute nodes with 48 cores and 1TB of RAM
- GPU nodes with Nvidia Tesla k40, Tesla k20, and Titan GPUs
- A high-speed Ethernet interconnect between 10 and 40 Gb/s

We used both CPUs and GPUs to train, evaluate, and predict statements of ignorance. We also used CPUs and GPUs for predicting annotations of textual mentions of OBO concepts.

Code for the ignorance-base and exploration methods can be found at: https://github.com/UCDenver-ccp/Ignorance-Base. The expanded ignorance corpus can be found at: https://github.com/UCDenver-ccp/Ignorance-Question-Corpus, with all associated code and models at: https://github.com/UCDenver-ccp/Ignorance-Question-Work-Full-Corpus. Code for concept recognition of the OBOs can be found at: https://github.com/UCDenver-ccp/Concept-Recognition-as-Translation.

### 3.2. Creating the ignorance-base

We created the ignorance-base grounded in knowledge goals and OBOs, extending prior work [20]. We expanded our corpus of ignorance statements based on knowledge goals to train and evaluate high-quality ignorance classifiers [4] and combined them with biomedical concept classifiers [25].

#### 3.2.1. Expanding the ignorance corpus

The goal was to create an ignorance corpus to show that ignorance statements can be reliably identified and automatically classified. We produced a gold-standard corpus consisting of articles with labeled sentences as **statements of ignorance** along with the **lexical cue(s)** (words or short phrases) that distinctly signify it as such mapped to a categorization of **knowledge goals** (**ignorance taxonomy**). This was done by examining spans of text each in the form of a word, short phrase, or whole sentence. Following the example above, “*<*these <u>inconsistent</u> <u>observations</u> point to the complicated <u>role</u> of VITAMIN D in the IMMUNE modulation and disease process*>*” (PMC4889866), the ignorance statement and entailed knowledge goal were identified based on the underlined words that communicate knowledge is missing, **lexical cues**, which map to an **ignorance taxonomy**, a formal categorization of knowledge goals. The cue <u>inconsistent</u> is an *indication of alternative research options or controversy of research* (abbreviated as *alternative/controversy*), and complicated is an *indication of difficult research task* (abbreviated as *difficult task*) (see Table 2). Our preliminary previous work [4] created a corpus of 60 articles annotated with lexical cues and ignorance categories. The goal was for an annotator to identify or an algorithm to classify that our example sentence was a statement of ignorance as shown by the brackets around the sentence. From there, once the sentence was deemed a statement of ignorance, the goal was to identify or classify that all underlined words including <u>inconsistent</u>, <u>observations</u>, etc. were the lexical cues that signified it as such. Note that one sentence can have multiple lexical cues that signify ignorance. The ignorance taxonomy helped to distinguish between different lexical cues: the annotator and classifier also needed to map the underlined cues to the specific ignorance category they deemed to capture the knowledge goal of the sentence. Here we expand that corpus to 91 articles to provide enough data to evaluate the classifiers on a held-out set of gold-standard data. We used the same methodologies as in our previous work [4], aside from a few minor changes.

Two new independent annotators, Katherine J Sullivan (K.J.S.) and Stephanie Araki (S.A.), both computational biology researchers, were provided with one to four articles, chosen randomly, in the Knowtator platform [147]. Each article was preprocessed such that lexical cues were automatically highlighted and linked to their corresponding classes of the ignorance taxonomy (Table 2), since prior efforts [4] showed that the annotation task was prohibitively difficult in unmarked documents. The annotators read through each article independently, deciding for each cue highlighted whether it signified an ignorance statement or not, and then either confirmed the ignorance taxonomy category or deleted the cue. Note that a lexical cue can map to multiple categories depending on the context (*e.g.*, the cue <u>however</u> can map to *anomalous/curious* or *alternative/controversy*). The annotators were also asked to add any new cues that signified ignorance and were not already highlighted by mapping them to the correct ignorance category. In the next annotation round, these new cues were added to the ignorance taxonomy. To capture the scope of each ignorance cue, we adapted the guidelines from BioScope [58] to highlight the whole sentence as the scope capturing all encompassed lexical cues due to difficulties with capturing only parts of the sentences. The annotators reviewed all annotations together, and Mayla R. Boguslav (M.R.B.) adjudicated any disagreements as they arose to create gold-standard articles that achieved an inter-annotator agreement (IAA) of at least 70-80%. The IAA is a measure of how well the annotations agree, and here we calculated the F1 score between the two annotations [150, 151]. An exact IAA was calculated on the exact text span of lexical cues or scopes as well as the ignorance category assignments. We also calculated a fuzzy IAA when the category assignments matched but not the text span of the cue or scope, or vice versa. For example, one annotator may highlight only <u>need</u> in a sentence containing the phrase <u>need to be</u>. In this case, we adopt the larger text span. (See [4] for more details).

K.J.S. and S.A. were trained first on eight random articles chosen from the 60 previous gold-standard articles. Any changes made to these articles (due to more experience with the task) were marked accordingly. After reaching the required IAA, new articles were chosen randomly using seeded randomness. For the first eight new articles (two batches of four), both annotators annotated the same articles as usual. After reaching IAAs of 80% or higher, we decided to divide the work: each annotator separately annotated one or two different articles and then adjudicated all annotations with M.R.B. Since the classic IAA could not be calculated because there was only one annotator, we calculated an “F1 score” between the original set of annotations and the adjudicated version to see how reliable the single annotation was compared to the final adjudicated version. We continued annotation when this score stayed above 80%, indicating a sufficient level of accuracy.

#### 3.2.2. Training and evaluating high-quality ignorance classifiers

With these extra articles, new classifiers were trained and optimized using a training set of 65 articles (approximately 2/3) and ultimately evaluated against a held-out test set of 26 articles (approximately 1/3). Ignorance classification can be made at either the sentence or the word level and as either a binary or a multi-classification problem. At the sentence level, the binary task determines whether or not a sentence’s scope contains a statement of ignorance. Since each statement of ignorance has at least one lexical cue labeled, the sentence can also be labeled using the ignorance categories implied by its lexical cues. Following the example above this would include the categories *alternative/controversy*, *difficult task*, etc. This now created a multi-classification problem of mapping sentences to the specific ignorance categories of their lexical cues. Conversely, we can also focus the binary task on the lexical cues to classify whether a word in an article was part of a lexical cue or not as labeled in the corpus. For the multi-classification task, the words would be mapped to their specific ignorance categories. Note that the test data included 501 unique lexical cues with no sentence examples in the training data. To avoid batch effects based on the different annotators, we split each batch separately (see Table 3).

**Table 3:** Data split for automatic classification: The table is in order of completion of annotation batches. Note that E.K.W. is Elizabeth K. White, M.R.B. is Mayla R. Boguslav, E.D. is Emily Dunn, Gold-standard is the previous gold-standard up to that point (the first row), K.J.S. is Katherine J. Sullivan, and S.A. is Stephanie Araki. *M.R.B. is an annotator along with the others. **E.D. only annotated one article along with the other annotators and then stopped. ***M.R.B. was the adjudicator in these batches.

| Annotation batch | Total Articles | Training Articles | Testing Articles |
| --- | --- | --- | --- |
| Prior work: E.K.W., M.R.B.*, (E.D.**) | 52 | 37 | 15 |
| Training: Gold-standard, K.J.S., S.A. | 8 | 6 | 2 |
| K.J.S., S.A., (M.R.B***) | 8 | 6 | 2 |
| K.J.S., M.R.B.*** | 11 | 8 | 3 |
| S.A., M.R.B.*** | 12 | 8 | 4 |
| Total Articles | 91 | 65 | 26 |
| Total Sentences | 12,055 | 8,281 | 3,774 |
| Total Words | 416,866 | 285,439 | 131,427 |

For both the sentenceand word-level tasks, we created both one true multi-classifier and also split each task up into 13 smaller binary tasks in which the ignorance category of interest was the positive case and all other sentences/words belonging to a different category were negative cases. Combining all 13 classifiers into an ensemble gave the full categorization for each sentence and avoided the problem of overlaps between categories. In all cases, we split the training data 90:10 for training and validation, and then evaluated separately on the held-out test set. We report the F1 scores for both sentence and word classification tasks on the held-out test set of 26 articles for (1) the one binary task (ALL CATEGORIES BINARY), (2) the 13 separate binary tasks that together create an ensemble multi-classifier (each category individually), and (3) the one multi-classifier (ALL CATEGORIES COMBINED - the macro-average).

Each article was segmented into sentences and then words used in the respective classification tasks. All models were chosen based on our prior work [4] and on an evaluation of several canonical models for concept recognition [25]. As our taxonomy was very similar to an ontology, we used our prior work in concept recognition [25] applied to a different type of linguistic phenomenon. In this work we explored and evaluated some of the canonical algorithms for concept recognition over many different ontologies and found that the CRF [152] and BioBERT [153] achieved the best performance for the task of span detection [25].

For sentence classification, we first built a simple Feed-Forward Artificial Neural Network (ANN), consisting only of three layers as our baseline model. We then fine-tuned both BERT [154] and BioBERT [153], so that we could compare a basic deep learning model to state-of-the-art language models. Our ANN consisted of a flattened layer followed by several dense layers to allow for arbitrary non-linear transformations of the input, with early stopping callbacks to avoid over-fitting (Additional details have been previously published [4]). For BioBERT, we used its domain-specific vocabulary to train the base BERT model. The same hyper-parameters were used for both BioBERT and BERT: batch size of 16, patience of 5, and a learning rate of 1 *×* 10^−5^. The number of epochs was tuned using truncating fuctions to avoid overfitting. We did not freeze the layers of the pre-trained BERT model and allowed the weights to keep updating during training for better performance.

For word classification, the goal was to automatically identify all lexical cues per our annotation task setup. We represented the underlying data using BIO-tagging: a word at the beginning of a lexical cue was marked ‘B’; a word inside a multi-word cue was marked ‘I’; a word outside of a cue (*i.e.*, not a word in the lexical cue) was marked ‘O’. If the lexical cue contained a discontinuity (*e.g.*, <u>no…exist</u> where the “…” signifies a discontinuity), we labeled the words that exist between the lexical cues as ‘O-’ (BIO-tagging scheme from [25]). CRF models were tuned with L1 and L2 regularization to avoid overfitting using the sklearn-crfsuite Python package [155]. For BioBERT, the named entity recognition baseline parameters performed quite well, most likely because it is a similar task, and so we did not tune any other parameters [153]. For a more thorough discussion of all data preparation and representation, the performance metrics, and the classification algorithms please refer to our prior work [4, 25].

Our ignorance statement identification task relied on the identification of lexical cues. To determine the role they played in classifying sentences, we conducted an ablation study. We deleted all ignorance-category annotations of the sentences and then re-trained the sentence classifiers using the best performing model for each category. We tested the models’ performance on our held-out test data set. A poor performance would indicate that the sentence classifiers rely heavily on lexical cues, while a good performance would point to the existence of other features beyond lexical cues that could identify ignorance statements. We report the results of these classifiers and discuss the use of lexical cues in related work to understand how well they apply beyond our work here. Additionally, we compared our cue list to some canonical work in the field to show generalizability beyond our prenatal nutrition corpus. We compared our list to the Bioscope [58] lexical cue list for clinical abstracts and articles, the lexical cue set for the meta-knowledge annotations of the GENIA project [156, 98], and the COVID-19 search engine keyword list [20]. We allowed for partial matching between the cues from each list to consider all forms that the cue can take within a phrase. Note that these works are similar to ours, but not the exact same task. Since each of these works focused on different domains than ours, the number of overlapping cues between our work and theirs may indicate generalizability beyond our work.

#### 3.2.3. Combining ignorance and biomedical concept classifiers

Creating an ignorance-base grounded in knowledge goals and OBOs allows us to explore it and provide analyses, summaries, and visualizations of the outputs. Further, grounding the ignorance-base in the OBOs allows us to connect our work to many other knowledge-bases [22]. To create the ignorance-base we combined the ignorance and biomedical concept classifiers over all 1,643 prenatal nutrition articles. The ignorance-base included all sentences from all articles to capture all biomedical concepts for comparison of our ignorance approach (only ignorance statements) to the standard literature search approach (all articles).

For ignorance classification, we used the 91 gold-standard corpus articles, and ran the best ignorance classifiers over the other 1,552 articles. Similarly, we ran our state-of-the-art biomedical concept classifiers [25] over all 1,643 articles to automatically identify biomedical concepts represented in ten OBOs, taking the best-performing models in terms of F1 scores, (CRFs [152] and BioBERT [153]). Most F1 scores ranged from 0.7-0.98 with the exception of PR at 0.53 (see Table 5 in [25]). Note that even though all of the classifiers performed close to the state of the art for the task at hand, they are still automated, so we draw conclusions cautiously. Because of this, we manually reviewed a random sampling of the identified biomedical concepts (a few hundred of each). The ten OBOs used for our work were the same used to manually annotate the CRAFT Corpus [157, 158], (a corpus of fulltext articles annotated along multiple syntactic and semantic axes, including extensive concept annotations):

1. Chemical Entities of Biological Interest (ChEBI)
2. Cell Ontology (CL)

a. Gene Ontology (GO): Gene Ontology Biological Process (GO BP)
b. Gene Ontology Cellular Component (GO CC)
c. Gene Ontology Molecular Function (GO MF)
3. Molecular Process Ontology (MOP)
4. NCBI Taxonomy (NCBITaxon)
5. Protein Ontology (PR)
6. Sequence Ontology (SO)
7. Uber-anatomy Ontology (UBERON)

For each of these ontologies, two sets of concept annotations were created for CRAFT (and appear in the public distribution): only proper classes of these OBOs and another adding in extension classes to better integrate the OBOs (created by the semantic annotation lead but defined in terms of proper OBO classes). We employed automatic concept recognition of our prenatal nutrition corpus with both the core OBOs and with the corresponding extended OBOs (suffixed with “ EXT”). Note that classification performance on the OBOs EXT was lower in general compared to the OBOs, especially for PR and PR EXT, so caution should be taken in interpreting those results. We focused on the proper OBO classes going forward, but have the data and results for both. (PheKnowLator does not currently have the OBOs EXT, but it is easily extendable.) Combining ignorance and biomedical concept classifiers automatically captured all ignorance statements and biomedical concepts for the 1,643 prenatal nutrition articles.

For clarification, the underlying data for the ignorance-base included sentences like the example above and another example: “it has an important <u>role</u> in BONE HOMEOSTASIS, BRAIN DEVELOPMENT and MODULATION OF the IMMUNE SYSTEM and yet the impact of ANTENATAL VITAMIN D deficiency on infant outcomes is poorly understood” (PMC4072587). The lexical cues important mapped to *important consideration*, <u>role</u> and impact mapped to *indefinite relationship*, yet mapped to *anomalous/curious*, and poorly understood to *unknown/novel*. BONE HOMEOSTASIS mapped to GO:0060348 (bone development), BRAIN DEVELOPMENT mapped to GO:0007420 (brain development), BRAIN also mapped to UBERON:0000955 (brain), MODULATION OF…IMMUNE SYSTEM mapped to GO:0002682 (regulation of immune system process), IMMUNE SYSTEM also mapped to UBERON:0002405 (immune system), ANTENATAL mapped to GO:0007567 (parturition), and VITAMIN D mapped to ChEBI:27300 (vitamin D). All of these mappings were identified by the classifiers. Note that we also identified biomedical concepts in non-ignorance statements. The entailed knowledge goal was to explore the relationship between prenatal vitamin D deficiency and infant outcomes through the important role of vitamin D.

The power of the ignorance-base is in its potential to be used for exploratory analyses, summaries, and visualizations to help researchers choose a topic to study or contextualize their experimental results in the known unknowns. Thus, in order to explore these data, we created a network representation of the ignorance-base to connect all sentences from these articles using both the ignorance lexical cues and biomedical concepts (see Figure 2). We combined all the literature data to connect sentences that have the same ignorance lexical cues, such as poorly understood, and then used PheKnowLator to compile all assertions mentioning the same given set of biomedical concepts, such as VITAMIN D. The semantic integration of PheKnowLator allowed us to not only connect our sentences to the biomedical concepts, but also to related ones; these connections were used in exploration by experimental results. This network can be used to search for all sentences that include the biomedical concept VITAMIN D, the lexical cue poorly understood, sentences with the ignorance category *unknown/novel*, or any combination of these features. Each sentence also related back to an article with its own metadata to be used for summaries and visualizations. For instance, the publication date was used to map how ignorance categories changed over time for a topic. Note that all sentences in all articles were included whether or not they contained ignorance statements, allowing for the ignorance enrichment comparison to the background information.

**Figure 2:**
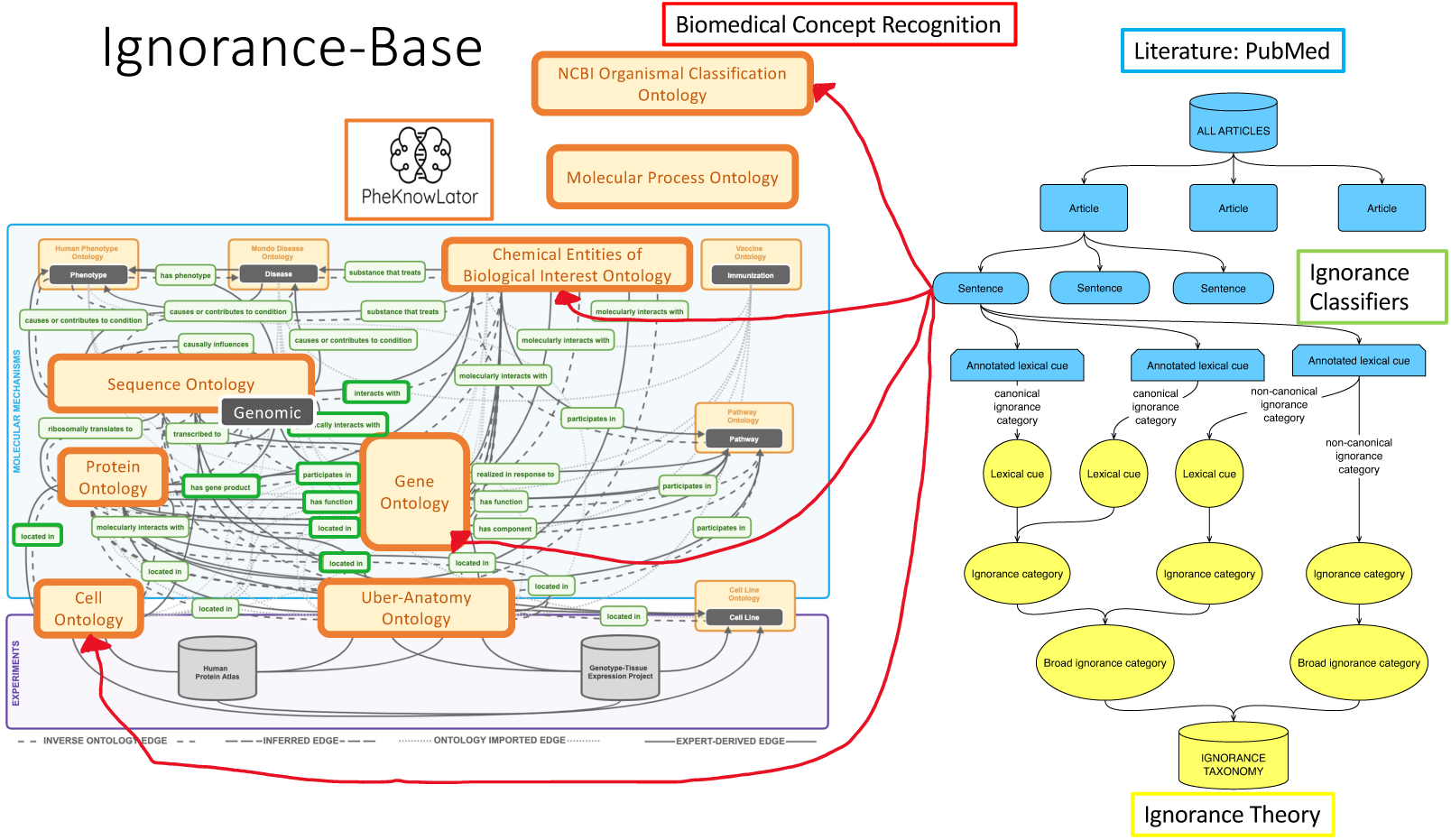
Network representation of the ignorance-base: The top right corner is the literature connecting the articles via segmented sentences (in blue) to the ignorance taxonomy (in yellow) through the ignorance classifiers (the annotated lexical cues). The sentences also connect to the biomedical concepts on the left with PheKnowLator [26, 27] using the biomedical concept classifiers with the ontologies of interest in bold and larger font.

### 3.3. Exploration by topic

The goal of exploration by topic was to explore the ignorance statements surrounding a topic to reveal novel insights. We compared our approach to a standard literature search (using biomedical concept expansion without ignorance expansion) and the sentences returned by the COVID-19 search engine [20]. Our comparison to the standard literature search provides a direct and informative comparison of results at the sentence level.

For our ignorance approach, an input topic consisted of a list of ontology concepts in PheKnowLator. To illustrate our approach, we explored the topic of vitamin D in consultation with a prenatal nutrition specialist (T.L.H.). We mapped the topic of vitamin D to four OBO concepts narrowed from 38 exact matches (280 partial matches): VITAMIN D (ChEBI:27300), D3 VITAMINS (ChEBI:73558), CALCIOL/VITAMIN D3 (ChEBI:28940), and VITAMIN D2 (ChEBI:28934). Note that going forward, when we refer to vitamin D, we mean the union of these four search terms. For the standard literature approach, we gathered all sentences from the ignorance-base that included terms from this vitamin D OBO concept list. For the ignorance approach, we only took the sentences that contained a vitamin D OBO list concept and had an ignorance lexical cue. For the COVID-19 search engine [20], we searched for the concepts “vitamin D” and “pregnancy” (to simulate our corpus theme), and compared their results to our approach. They provided a list of sentences and a drop-down list of the most frequent concepts in relation to the topic.

For our ignorance approach, we not only provided a list of sentences and the most frequent concepts, but also an analysis of the most ignorance-enriched concepts (ignorance enrichment). The goal was to help researchers narrow in on a specific topic to explore in future research. The ignorance statements surrounding the topic were available to explore. We also provided visualizations of the most frequent concepts: we present both biomedical concept clouds and word clouds along with frequency tables to explore them. For enrichment, we used the hypergeometric test for over-representation with both Bonferroni (family-wise error rate) and Benjamini-Hochberg (false discovery rate) multiple testing corrections [159, 125] to find concepts enriched in ignorance statements compared to all vitamin D sentences. We compared these results to the standard literature search approach using concept enrichment (concepts enriched in all vitamin D sentences compared to all sentences). In comparing our ignorance approach to the standard literature approach (see Figure 3), if a concept was more frequent or enriched in the standard approach but not in ignorance, then it could be established information. If a concept appeared in both approaches, then it might currently be under study (currently studied). If a concept only appeared in the ignorance approach then it could be an emerging topic. Note that concepts that were not frequent or enriched in either approach were not of interest. For the COVID-19 search engine [20], we could only compare the frequency lists.

**Figure 3:**
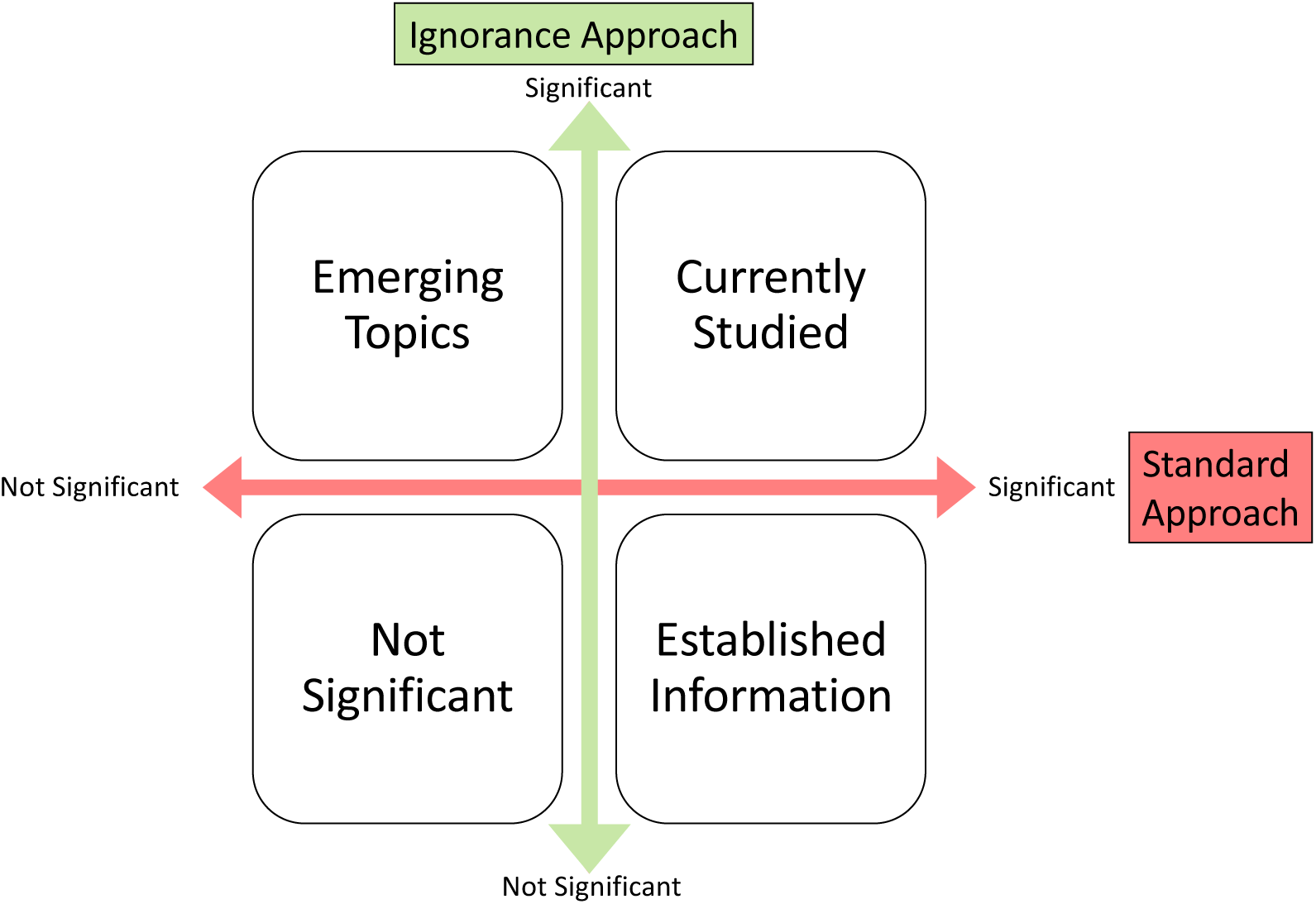
Ignorance vs. Standard Approach Results Chart: The interpretation of the results comparing the ignorance approach to the standard approach.

Our ignorance approach can further help researchers narrow their research topic to one with the right scope of ignorance. We demonstrate this by visualizing how known unknowns are described (ignorance-category enrichment) and how they change over time. We bubble plotted the ignorance categories per article over time, with the bubble size representing the percentage of sentences in an article scaled by the total number of sentences of that category. Using these methods, the researchers continued to deep dive into ignorance statements that included their topic, enriched concepts, and enriched ignorance categories to find knowledge goals to pursue for research. In order to determine if we found *novel* avenues to explore using our ignorance approach, we consulted both our prenatal nutrition specialist (T.L.H.) and PubMed (after our data collection) to corroborate our findings.

### 3.4. Exploration by experimental results

The goal of exploration by experimental results was to identify questions (ignorance statements) that may bear on the results, providing new avenues for exploration (biomedical concepts), potentially from other fields. Exploration by experimental results used the same methods as exploration by topic with some added pre-processing steps and analyses based on the relationship between the inputs. As before, the input topic was still an OBO concept list, but an extra pre-processing step connected the experimental results to OBO concepts in PheKnowLator [26, 27]. In general, as long as the experimental results can be mapped to OBO concepts in and through PheKnowLator, we can connect them to the ignorance-base.

To illustrate our approach, we used our motivating example, the vitamin D and sPTB gene list (Entrez genes) [36], and mapped it to the genomic part of the sequence ontology (SO) and the corresponding proteins in the protein ontology (PR). This initialized the list of ontology terms to use for our search. To add more terms, we utilized the relations ontology (RO) which connects the different ontologies together in PheKnowLator. For instance, the relation “interacts with” (RO:0002434) connects proteins or genes to chemicals (ChEBI). This yielded a large list of ontology terms; we then found all the sentences that contained these terms (our sentences of interest). Note that not all OBO concepts connected to a sentence. From here, we performed all the same analyses as exploration by topic, including finding articles, sentences, ignorance categories, and concepts to investigate. Further, we added three more analyses: (1) gene list coverage (prioritizing the OBO concepts that connect to the most genes), (2) comparisons to other enrichment analyses such as DAVID, and (3) comparisons to any other findings about the gene list such as findings from a paper. (See figure 4 for the exploration by experimental results pipeline.) Note that neither a standard literature search nor the COVID-19 search engine can currently support queries by experimental results.

**Figure 4:**
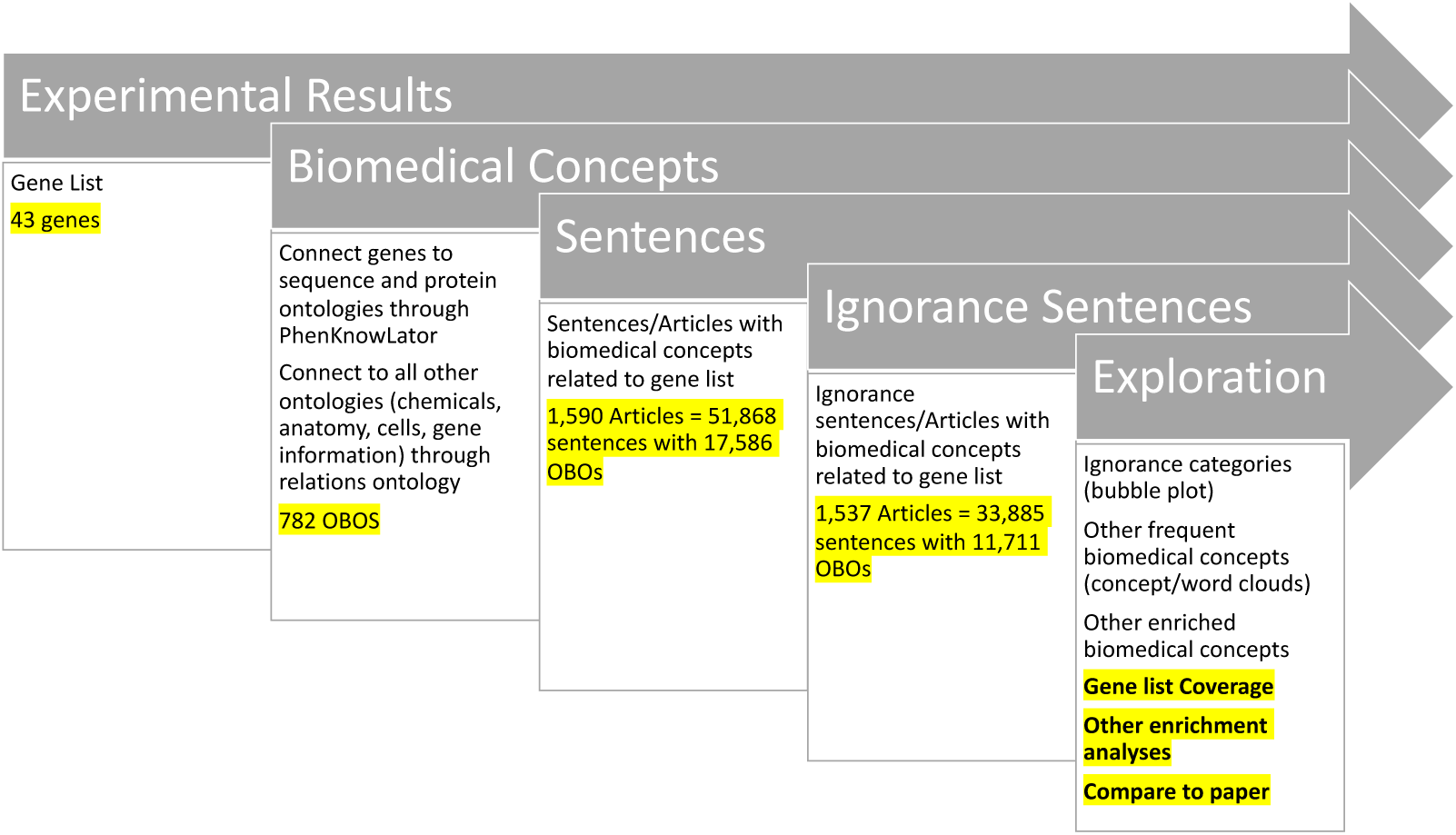
Exploration by Experimental Results (gene list) pipeline: The results are in yellow highlights for the example presented here. For exploration at the end of the pipeline, the three not highlighted are the same as exploration by topic and the three highlighted are the new additions based on a gene list.

Gene list coverage can help prioritize which OBO concepts are most critical to examine. As we mapped the gene list to the OBO concepts, some OBO concepts had many genes map to them, implying that these OBO concepts were potentially more relevant to the gene list than concepts with fewer genes mapping to them. Thus, we sorted the OBO concept list by these high coverage ones and looked to see if those were enriched in all of our sentences of interest and/or in ignorance sentences. This provided a smaller and more refined list to start exploring.

Since canonical enrichment methods can also help prioritize OBO concepts, we compared our ignorance enrichment method to them, allowing us to both enhance the canonical methods and find new lines of investigation. From tools such as DAVID [145], we got a list of enriched OBOs (GO concepts) by using the gene list and functional annotations from their entailed knowledge-bases. We then found and examined any ignorance statements that contained the concepts linked by DAVID. Further, our method provided a list of OBO concepts enriched in ignorance statements. Thus, we compared and examined these two lists to add the ignorance layer to the classic enrichment analysis and to find new concepts that may be currently unknown to the knowledge-bases but potential emerging topics related to the gene list.

Given that our gene list came from a paper, we compared our ignorance-approach findings to the paper findings to identify questions that may bear on it, providing new avenues for exploration from other fields. Yadama *et al.*, [36] focused on the immune system in their main findings, but we also found a new path in brain development for the researchers to explore, based on concepts and knowledge goal statements not mentioned or cited in the article. We also found later review articles that corroborate this novel direction.

## 4. Results

### 4.1. The ignorance-base: The power of combining ignorance and biomedical concept classifiers

The ignorance-base captured the connection between our collective scientific ignorance (ignorance taxonomy) and knowledge (PheKnowLator) through sentences from the literature and yielded a wealth of data (see Figure 5) for future study via the network (see Figure 2). The short manual review of some random sentences from the ignorance-base suggested that both the ignorance and biomedical concept classifiers correctly identified concepts (data not shown). Combining these two types of classifiers enhanced the exploration methods.

**Figure 5:**
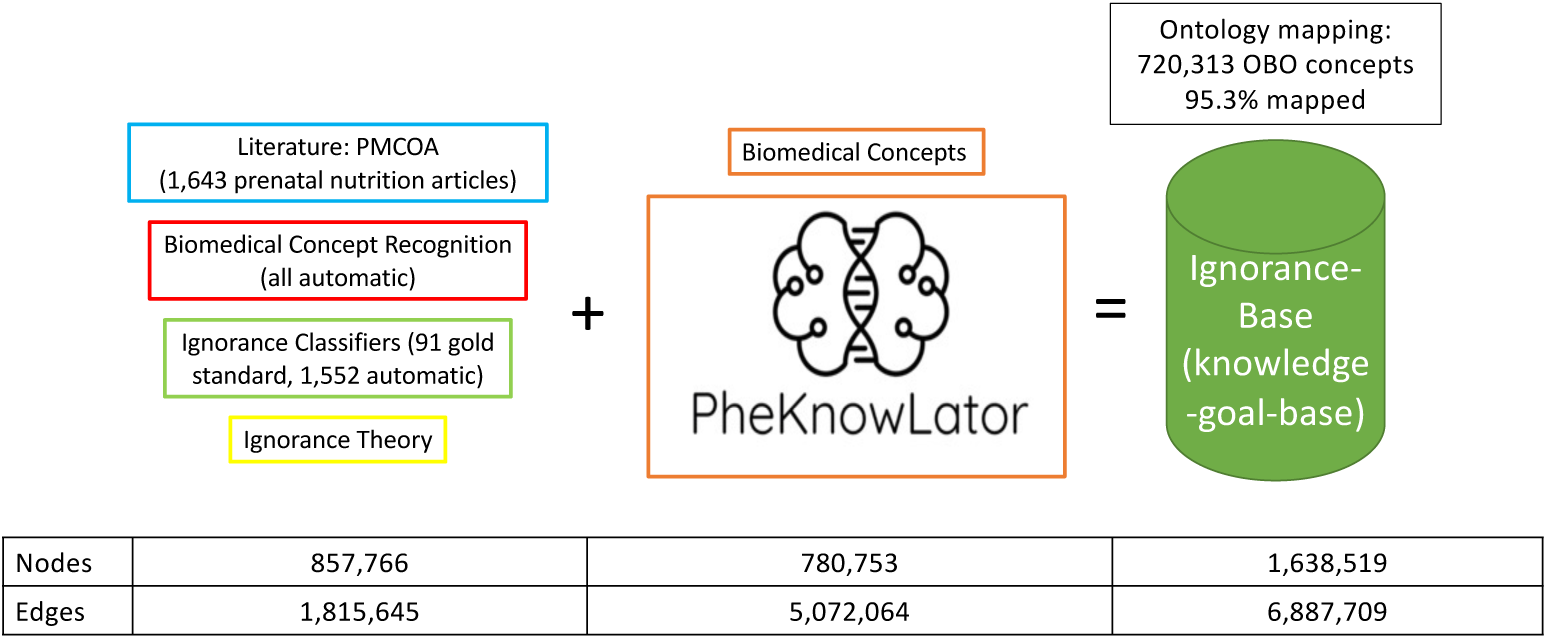
Summary information for the ignorance-base. The ignorance-base is a combination of biomedical concept classifiers and ignorance classifiers over a corpus of prenatal nutrition articles. The network representation connected the literature to the ignorance theory and biomedical concepts via PhenKnowLator [26, 27].

Creating an expanded gold-standard ignorance corpus (see Table 4) yielded ignorance classifiers achieving F1 scores around or above 0.8 with many closer to 0.9, on both the sentence-and word-levels (see Tables 5 and 6). The ensemble of 13 different binary classifiers performed the best for both classification tasks and was used for all ignorance classification for the ignorance-base. These high-quality classifiers, built on the expanded ignorance corpus, allowed us to scale up our system for the ignorance-base.

**Table 4:** Interannotator Agreement (IAA): IAA is calculated as F1 score for all annotation tasks. The IAA for the training is between the two annotators, not including the previous gold-standard. *F1 score between annotator and the final gold-standard version after adjudication with M.R.B.

| Annotation batch | category IAA | scope IAA | fuzzy category IAA | fuzzy scope IAA |
| --- | --- | --- | --- | --- |
| Prior work (60 articles) | 78% | 87% | 79% | 90% |
| Training (8 articles) | 77% | 66% | 78% | 87% |
| K.J.S. and S.A. (8 articles) | 81% | 82% | 81% | 93% |
| K.J.S. with M.R.B adjudicator* (12 articles) | 88% | 92% | 89% | 95% |
| S.A with M.R.B adjudicator* (12 articles) | 89% | 92% | 90% | 96% |
| All combined | 82% | 87% | 83% | 92% |

**Table 5:**
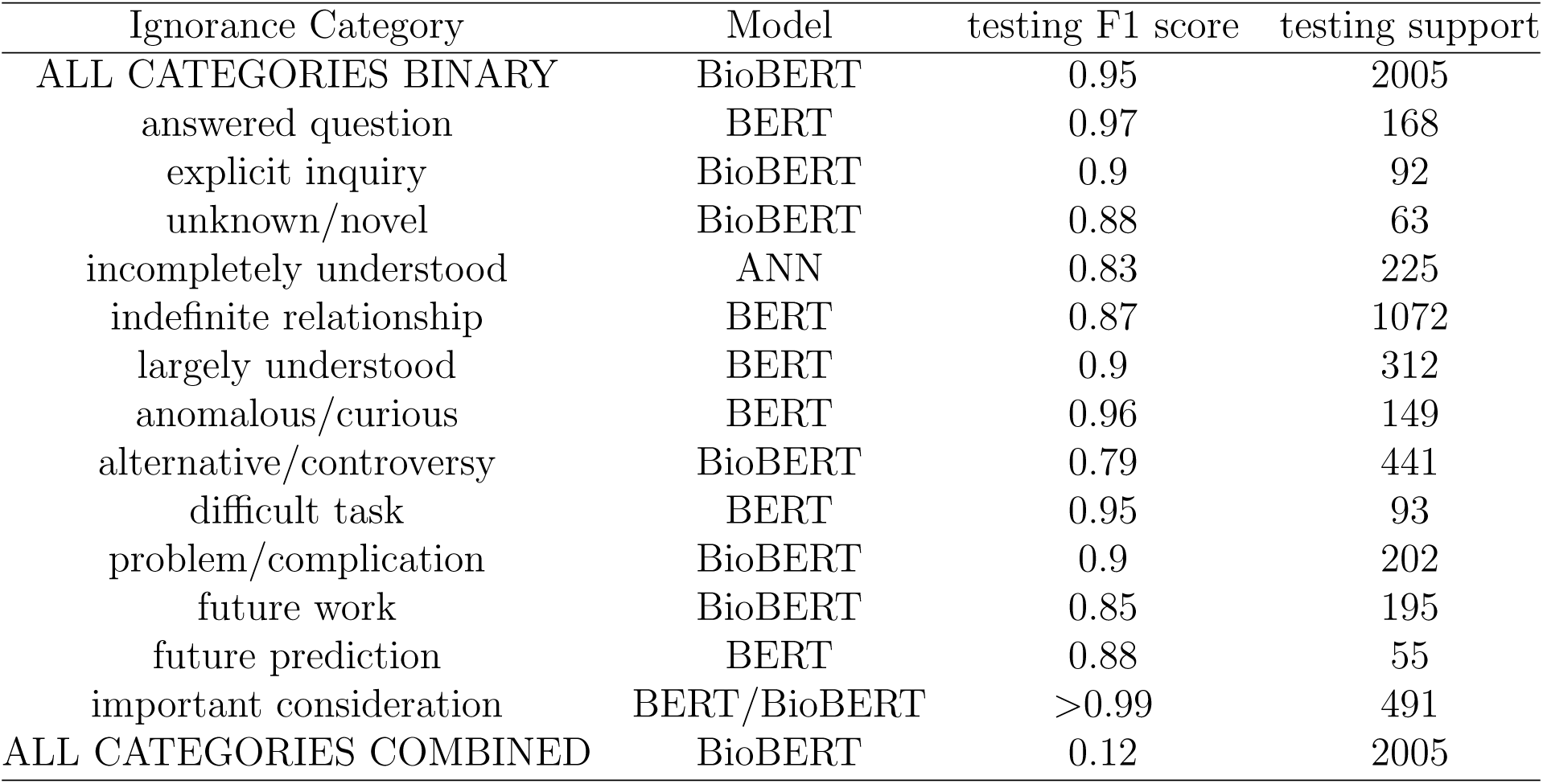
Sentence Classification: the best model for sentence classification for each ignorance category and all categories combined.

**Table 6:** Word Classification: the best model for word classification for each ignorance category and all categories combined. *Reporting the average F1 score of all the categories for one multi-classifier.

| Ignorance Category | Model | testing F1 score | testing support |
| --- | --- | --- | --- |
| ALL CATEGORIES BINARY | BioBERT | 0.89 | 7601 |
| answered question | BioBERT | 0.89 | 320 |
| unknown/novel | CRF | 0.98 | 155 |
| explicit inquiry | BioBERT | 0.97 | 43 |
| incompletely understood | BioBERT | 0.93 | 2809 |
| indefinite relationship | BioBERT | 0.97 | 1205 |
| largely understood | BioBERT | 0.94 | 618 |
| anomalous/curious | BioBERT | 0.96 | 399 |
| alternative/controversy | BioBERT | 0.91 | 598 |
| difficult | CRF | 0.93 | 128 |
| problem/complication | BioBERT | 0.9 | 238 |
| future work | BioBERT | 0.89 | 391 |
| future prediction | BioBERT | 0.94 | 100 |
| important consideration | BioBERT | 0.93 | 608 |
| ALL CATEGORIES COMBINED* | BioBERT | 0.82 | 6239 |

For the corpus, the annotation guidelines may be generalizable, with five different annotators over all annotation tasks. We can trust the annotations and reliability of the guidelines as the IAA, as measured by an F1 score, was near or above 80% for the classic annotation task [150, 151, 160] and for the split annotations, the annotators were correctly identifying statements of ignorance around 90% of the time. The task was quite difficult, requiring the pre-processing of the documents, extensive training, and many examples of ignorance statements. Disagreements involved the annotators choosing different lexical cues that signified ignorance for a given sentence, different semantic interpretations of a sentence, and the need for clarification on the ignorance categories. Consensus was reached during the adjudication process and the annotation guidelines and ignorance taxonomy were updated based on those discussions. Our annotation guidelines were robust and reproducible in two different annotation tasks. (For more information on the corpus itself see the Supplementary File on Corpus Information.)

Our results suggest that ignorance statements proliferate throughout the ignorance-base. The 1,643 articles, spanning years 1939 to 2018 (see Figure 6), contained 327,724 sentences with over 11 million words. Just over half of those sentences had an ignorance lexical cue (182,892), with articles averaging a total of 111 cues (with a median of 93). Every section of the articles had ignorance cues aside from the title, with the most in the discussion and conclusion sections and the fewest in the abstract and results.

**Figure 6:**
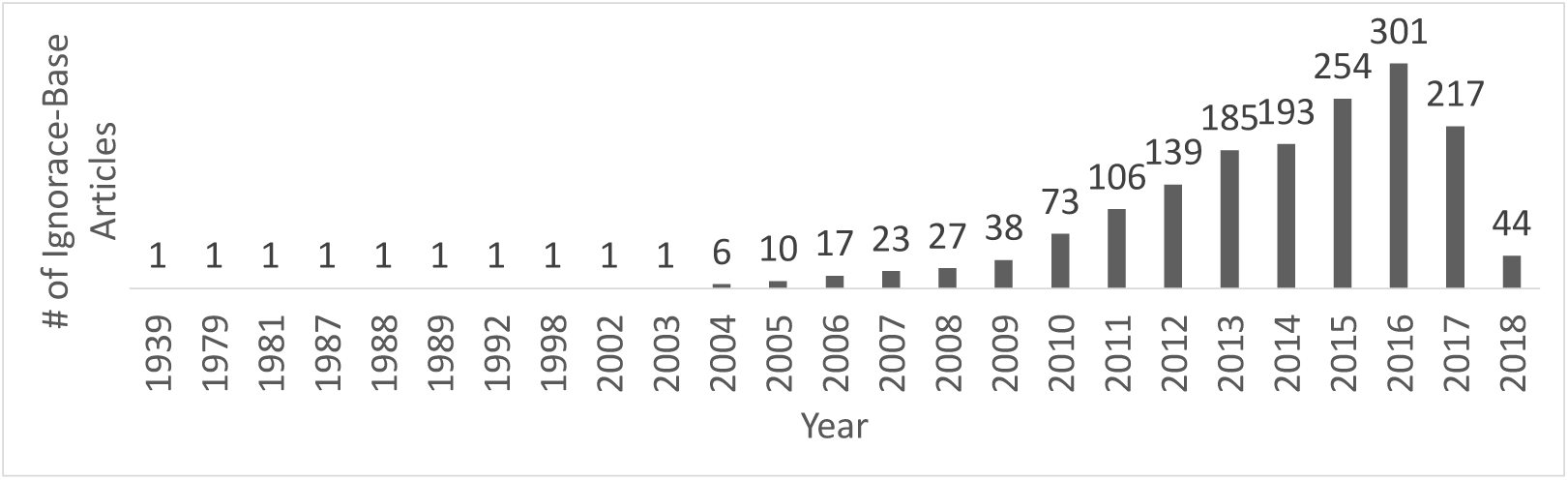
Article date distribution for the ignorance-base (1939-2018).

Our collective scientific ignorance is abundantly represented throughout the literature.

Further, our ignorance taxonomy is not only a categorization system of ignorance statements via knowledge goals, but also a depiction of the research life-cycle and how researchers discuss our collective scientific ignorance (see Figure 7 with proper definitions and example lexical cues in Table 2). (Note that we renamed our ignorance categories from our previous work [4] to be more ontologically precise based on discussions with an ontologist, Mike Bada.) Underneath the 13 categories of ignorance (with 3 broader ones in all caps in Figure 7) were 2,513 unique lexical cues collected from related work or added during our annotation tasks. 1,822 of them had examples in our ignorance-base. Further, the ignorance classifiers found 5,637 new unique lexical cues that signify ignorance. These new cues were added to the ignorance-base and noted as such. Many of these cues were variations of ones already captured and others were new, such as “not as yet”, “interplay”, and “have begun to illuminate”. Our ignorance classifiers recognized more complex language than just a dictionary match. In addition, the classifiers quite heavily relied on the lexical cues as features (see Table 7). Our ablation study showed poor performance without them. We found that the performance on the held-out test set dropped significantly for all ignorance categories, with an average F1 score of 0.39. Further, our cues seem to generalize beyond prenatal nutrition and our corpus based on the many overlaps between our cue list and prior work from different biomedical domains (see Table 8). Overall, there were 517,445 ignorance annotations involving 7,459 unique lexical cues; this reinforces the diversity of ways that ignorance is expressed in the literature.

**Figure 7:**
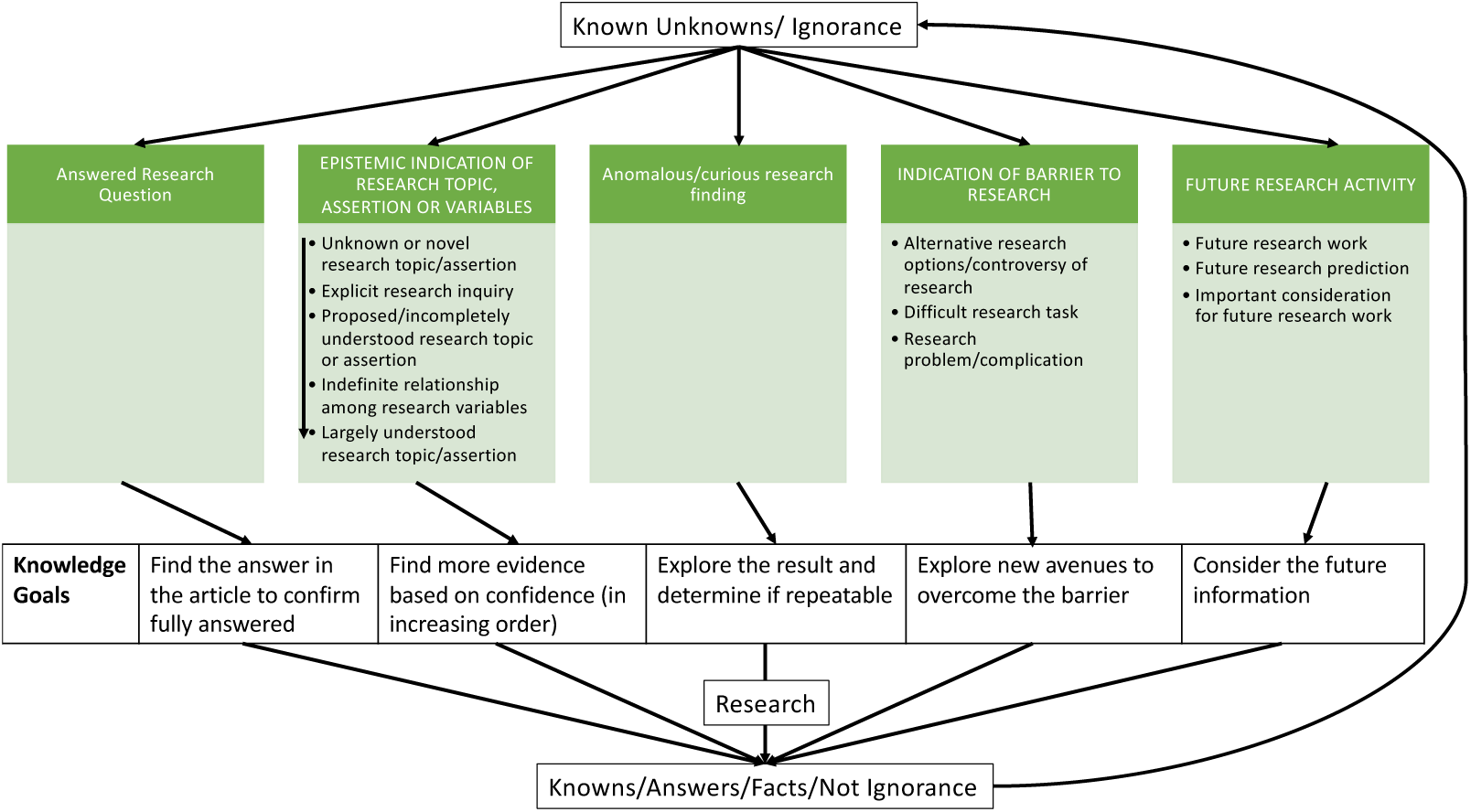
Ignorance taxonomy embedded in the research context: Starting from the top, research starts from known unknowns or ignorance. Our ignorance taxonomy is in green (an ignorance statement is an indication of each ignorance category) with knowledge goals underneath. Research is then conducted based on the knowledge goals to get answers; these then filter back to the known unknowns to identify the next research questions.

**Table 7:** Ablation study: Results of the ablation study on the sentence classification level.

| Ignorance Category | Model | testing F1 score | testing support |
| --- | --- | --- | --- |
| ALL CATEGORIES BINARY | BERT | 0.33 | 2005 |
| answered question | BERT | 0.47 | 168 |
| explicit inquiry | BERT/BioBERT | 0.35 | 92 |
| unknown/novel | BERT/BioBERT | 0.25 | 63 |
| incompletely understood | BERT | 0.27 | 225 |
| indefinite relationship | BERT | 0.52 | 1072 |
| largely understood | BERT | 0.49 | 312 |
| anomalous/curious | BERT | 0.58 | 149 |
| alternative/controversy | BERT | 0.23 | 441 |
| difficult task | BERT | 0.35 | 93 |
| problem/complication | BioBERT | 0.43 | 202 |
| future work | BioBERT | 0.61 | 195 |
| future prediction | BERT | 0.22 | 55 |
| important consideration | BERT/BioBERT | 0.46 | 491 |

**Table 8:** Lexical cue generalizability study: Results of overlaps between our lexical cue list (2,513 cues) and other similar works from different domains.

| Similar work | cues support | matching support |
| --- | --- | --- |
| BioScope corpus [58] | 46 | 45 (98%) |
| GENIA Project (Meta-knowledge) [156, 98] | 1374 | 591 (43%) |
| COVID-19 Search Engine [20] | 286 | 166 (60%) |

Our ignorance-base also contained a wealth of different types of biomedical concepts. Our biomedical concept classifiers identified 720,313 concepts involving 19,883 unique concepts from all of the ontologies and almost all of them (95.3%) mapped to PheKnowLator. Note that we can only represent the biomedical concepts in PheKnowLator that were also captured by our biomedical concept classifiers. This overlap included six of our eight ontologies (missing MOP and NCBITaxon) or six of the eleven PheKnowLator ones (missing the human phenotype ontology, MONDO disease ontology, vaccine ontology, pathway ontology, and cell line ontology). Because our biomedical concept classifiers predict identifiers character by character (see our prior work for more details [25]), they can produce identifiers that do not exist. In terms of errors, our classifiers predicted 850 (4%) unique OBO concepts with non-existent OBO identifiers. The other 1,432 (7%) unique OBO concepts that did not map seemed to either be from the two ontologies not included in PheKnowLator (MOP and NCBITaxon) or were terms no longer used/depreciated from the ontologies. The ignorance-base captured many biomedical concepts. Overall, the ignorance-base contained a great deal of data consisting of all different types of lexical cues, ignorance categories, and OBO concepts.

### 4.2. Focusing on ignorance statements provides an alternative targeted exploration of a topic that is distinct from the standard approach

Focusing on ignorance statements provided researchers interested in the topic of vitamin D with new avenues of exploration that are distinct from the standard literature approach and the COVID-19 search engine [20] (see Figure 8). We present results for the ignorance approach (vitamin D ignorance statements), the COVID-19 search engine [20], the standard approach (vitamin D sentences only), and a comparison of all three.

**Figure 8:**
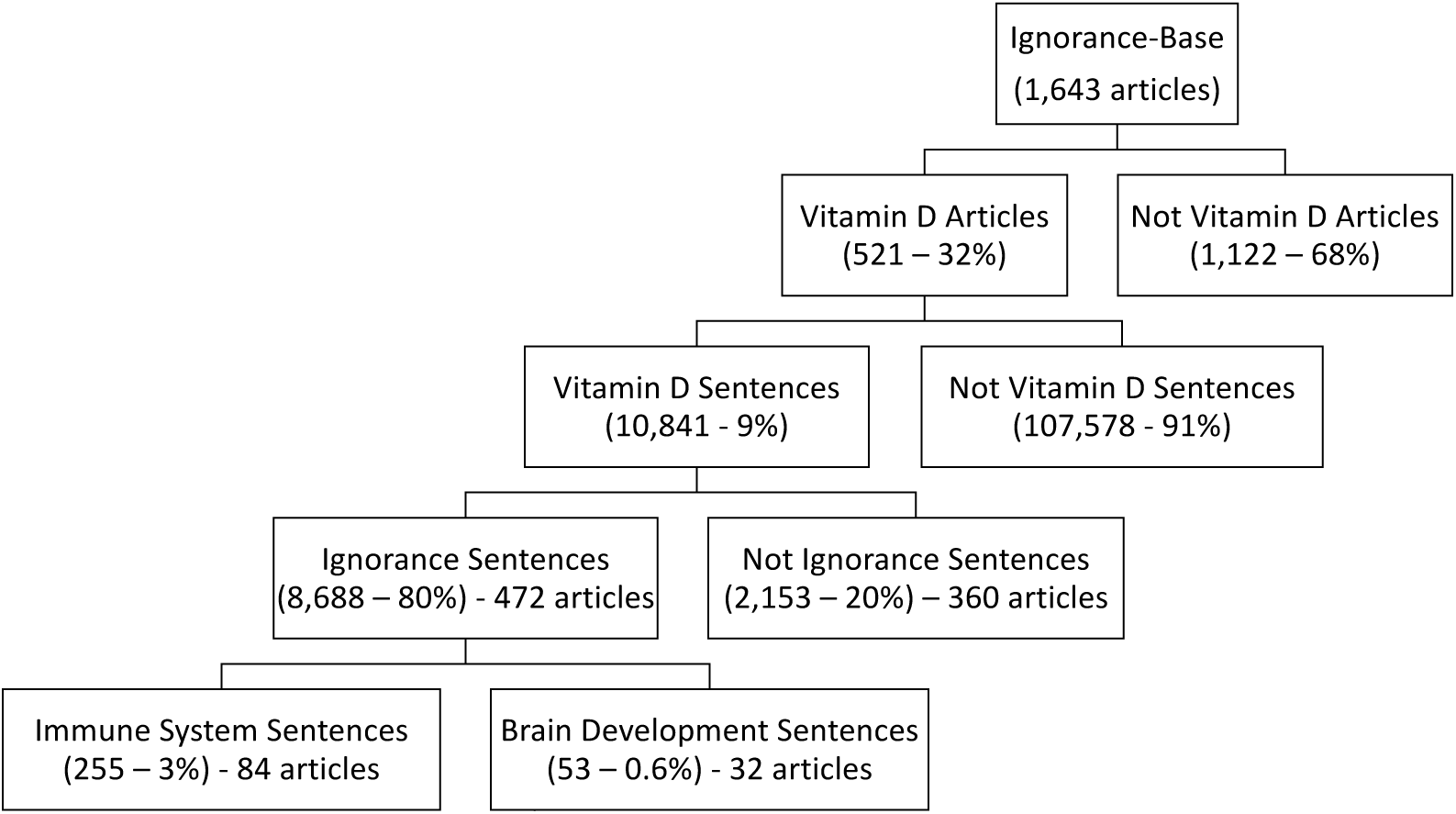
Exploring the ignorance-base by Vitamin D: Searching the ignorance-base for vitamin D yields many articles and sentences that can be explored using ignorance statements to find new research questions, including immune system and brain development.

There is a great deal of research on vitamin D and more specifically a plethora of ignorance statements. Searching through the ignorance-base for the four vitamin D terms yielded 521 articles with 10,841 sentences mentioning vitamin D (9% of the 118,419 sentences in the 521 articles and 3% of all the sentences in the ignorance-base) (see Figure 8). Note that only the terms VITAMIN D and VITAMIN D2 pulled out sentences from the ignorance-base. These sentences included 17,584 unique biomedical concepts excluding the VITAMIN D concepts (88% of the total unique biomedical concepts). Of those VITAMIN D sentences, 8,688 sentences (80%) were ignorance statements spanning 472 articles. We explored this data to differentiate between knowns and unknowns.

#### 4.2.1. Term Frequency

Focusing on term frequency provided some concepts of interest. The top five most frequent biomedical concepts for the ignorance approach included: FEMALE PREGNANCY, PARTURITION (giving birth), VERSICONOL ACETATE, BLOOD SERUM, and FEEDING BEHAVIOR (see Figure 9a). The first two aligned with the corpus theme of prenatal nutrition. VERSICONOL ACETATE is an intermediate in the biosynthesis of aflatoxin, a toxin produced by mold that may be toxic towards the vitamin D receptor in relation to rickets [161]. Vitamin D levels are mainly measured from the BLOOD SERUM, and FEEDING BEHAVIOR seems to highlight the importance of ingesting vitamin D. For the words, the most frequent terms were supplementation, maternal, status, levels, and women and they also fit with the theme: supplements are suggested for many people, maternal and women fit with the corpus theme, and status and levels are measurement terms for vitamin D. None of these terms were surprising, which was a good sign that we captured meaningful information.

**Figure 9:**
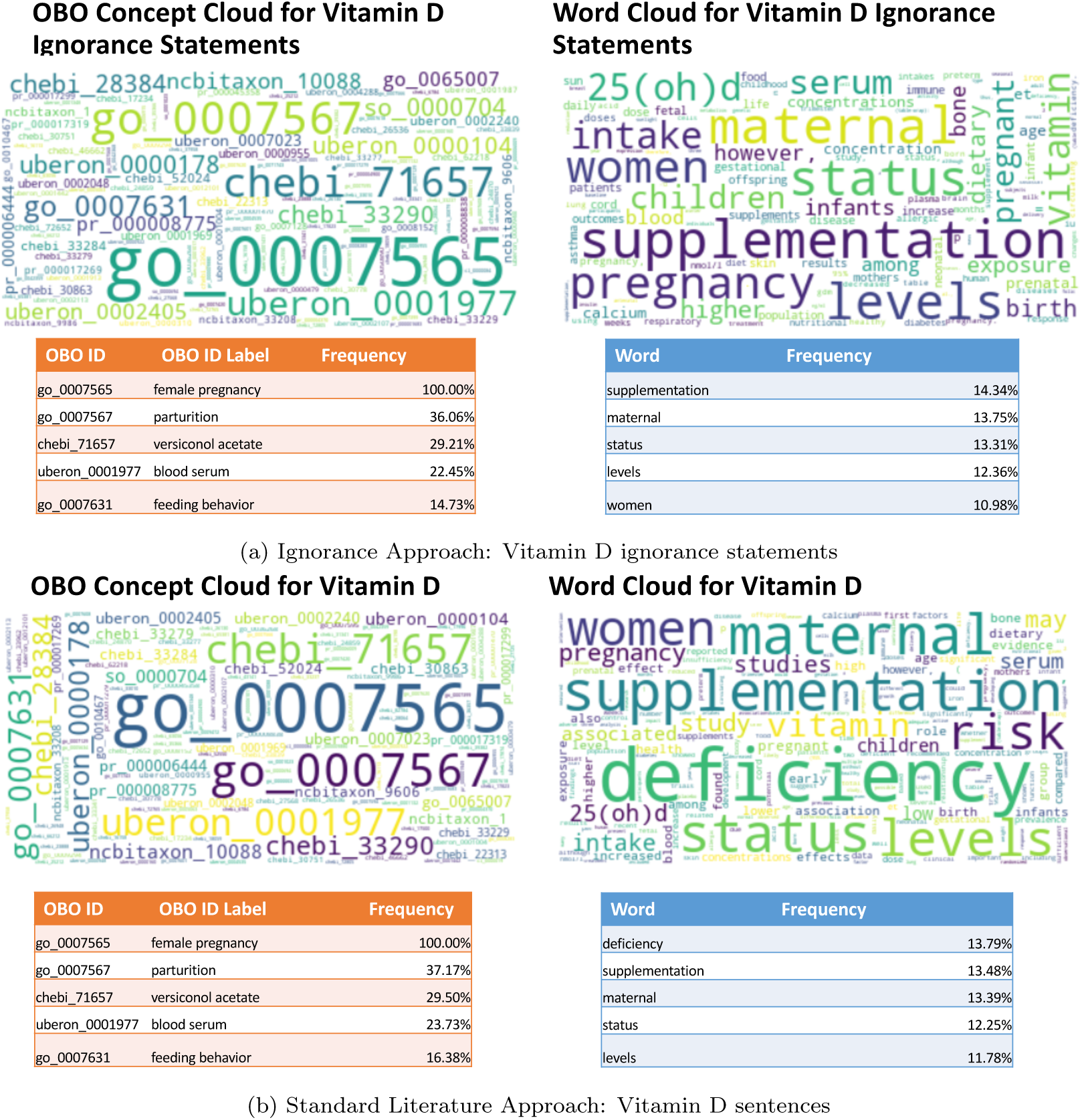
Term frequency results: Frequent Biomedical Concepts and Words in (a) ignorance approach vitamin D ignorance statements and (b) standard literature approach vitamin D sentences. Word clouds using words and biomedical concepts are on the right and left respectively. Also underneath are frequency tables of the top 5 most frequent concepts or words.

Term frequency can help prioritize areas to explore. Taking one of these terms or concepts provided avenues for the researchers to explore. For example, FEEDING BEHAVIOR (GO:0007631), defined as the behavior associated with the intake of food, was an interesting concept in relation to vitamin D. Vitamin D is naturally absorbed through sunlight and digestion. To corroborate our findings, we searched for “vitamin D” and “feeding behavior” in the literature, and found that vitamin intake during pregnancy in general seems to affect both the metabolic system and food intake regulatory pathways in the offspring [162]. This result argues that concepts that appear frequently with a topic can provide useful keyword search terms for researchers.

Further, frequent concepts can lead to some pertinent questions for the researchers. The ignorance statements for FEEDING BEHAVIOR and VITAMIN D, provided research ideas. Most of these ignorance statements discussed the ingestion of vitamin D mainly via supplements, but also with some foods. The recommendations for ingestion all varied by study (agreeing with the findings from a systematic review [163]). One ignorance statement stood out specifically, “the high prevalence of Vitamin D deficiency in PREGNANT women is a worldwide health problem regardless of latitude, FOOD INTAKE or socio-economic status [93]” (PMC5941617) [164], citing a systematic review and meta-analysis that looked at vitamin D status globally [165]. All of these studies recommended vitamin D supplementation, but we could not find any studies that determine why supplementation is so low. How do supplements, specifically for vitamin D, fit into feeding behavior? A potential research topic could be to study what specific factors, beyond general socio-cultural factors, lead to women taking vitamin D supplements as part of their diet, especially for pregnancy. Studying this topic could lead to novel methods that help mothers stay vitamin D sufficient throughout pregnancy, resulting in fewer adverse outcomes for both mother and offspring (see Figure 1). Thus, biomedical concept frequency can lead to a high impact research topic that could affect mothers of childbearing age and their offspring globally.

Term frequency can also highlight when the context of a term is more known or unknown. Comparing the ignorance approach to the standard literature approach, the most frequent concepts in the standard approach were the same as the ignorance approach (see Figure 9b). This suggests that that term frequency may not capture the difference in biomedical subjects between the two approaches. However, the top five words slightly differed between them: the word “deficiency” was the top most frequent term in the standard approach and the term “women” disappeared (see Figure 9). This may signify that vitamin D deficiency was established information, resulting in a lack of ignorance. At the same time, all these terms may be more unknown than known. Recall that 80% of the vitamin D sentences were ignorance statements, so it is possible that much of the context around VITAMIN D was still unknown in general. The ignorance frequency term list not only provided an avenue for exploration, FEEDING BEHAVIOR, with a potential research topic, but in addition, may help distinguish between terms that describe probable knowns, like “deficiency”, and those connected to more open questions, like FEEDING BEHAVIOR.

Comparing these ignorance results to the COVID-19 search engine [20] provided a different set of top frequent terms as MeSH terms. Searching for “vitamin D” and “pregnancy” in the COVID-19 search engine [20] resulted in 26 sentences. Within those sentences, the five most frequent terms were vitamin D deficiency, asthma, autoimmune diseases, child, and placenta with only 2-3 sentences for each of them. Note that the small number of sentences was probably due to the differing underlying themes of the corpora (COVID-19 mainly vs. prenatal nutrition). In comparing all frequency lists, we found the word “deficiency” in the standard approach with almost 14% of the corpus containing it. The terms “child” and “placenta” fit the theme of pregnancy and were similar to the terms found in the ignorance approach. The terms “asthma” and “autoimmune disease” raised possible avenues to explore, but were underrepresented with only two sentences per topic. Our ignorance approach provided more sentences, visualizations, and avenues to explore for researchers interested in prenatal nutrition topics including vitamin D.

#### 4.2.2. Ignorance Enrichment

To go beyond term frequency and further distinguish between known and known unknowns, ignorance enrichment found at least three interesting new avenues to explore in relation to vitamin D that were captured by the standard approach, but buried amongst 275 concepts, and one avenue not captured by the standard approach at all. Note that the COVID-19 search engine [20] did not calculate enrichment and so we can only compare to their frequency list. The ignorance approach found 11 ignorance enriched concepts, whereas the standard approach found 275, with an overlap of eight concepts (see Figure 10). However, only focusing on the overlapping concepts, in the standard approach most of them were buried far down the list of enriched concepts ordered by enrichment p-value (indicated by the parentheses next to the overlapped concepts in Figure 10). Further, in comparing the two different approaches, the ignorance approach found concepts from broader categories, including IMMUNE SYSTEM and BRAIN DEVELOPMENT, compared to the standard approach which were more specific entities, such as BLOOD SERUM and VITAMIN K. Ignorance enrichment provided the researchers with a smaller list of targeted statements of knowledge goals to potentially pursue or spark ideas from.

**Figure 10:**
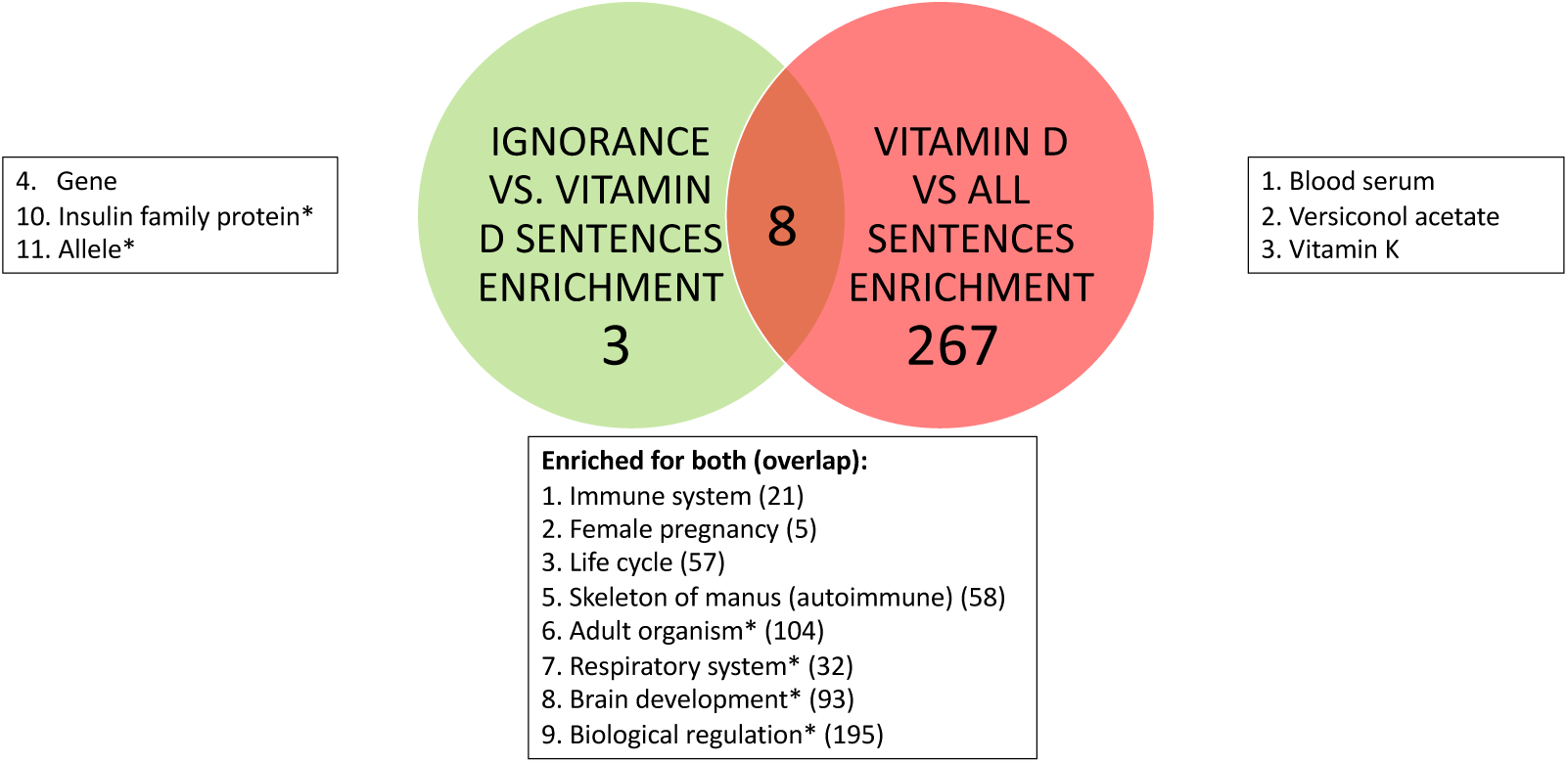
Comparison of standard and ignorance enrichment: A Venn diagram of biomedical concept enrichment between just vitamin D (pink) and ignorance vitamin D (green) sentences. Next to each bubble are concepts in their respective enrichment orders. The concepts in the middle are the overlap and the numbers correspond to the enrichment position for the ignorance vitamin D enrichment, with the overlap position in parentheses. Skeleton of manus is an error and is actually annotating autoimmune as in the parentheses. *Statistically significant with FDR but not family-wise error.

Focusing on the ignorance-enriched concepts also provided research topic ideas. T.L.H. and M.R.B. determined that IMMUNE SYSTEM, RESPIRATORY SYSTEM, and BRAIN DEVELOPMENT (all captured by the standard approach) were all interesting in relation to vitamin D. Intriguingly, the COVID-19 search engine frequency list included the terms “autoimmune diseases” and “asthma”, which are subsets of IMMUNE SYSTEM and RESPIRATORY SYSTEM, respectively. All of these concepts were currently studied, with more room for future work. We also found the insulin family protein (not captured by the standard approach) intriguing because many studies have attempted to determine the link between vitamin D and gestational diabetes mellitus [166]. All of these concept areas are ripe for exploration.

We explored the specific ignorance statements for the concepts of interest to narrow in on a research topic, just as with FEEDING BEHAVIOR. We chose to look at the IMMUNE SYSTEM ignorance statements, which provided the researcher with 255 ignorance statements plus their entailed knowledge goals, spanning 84 articles (see Table 9 for the top eight articles with the most ignorance statements). Note that only one article had no ignorance statements that included VITAMIN D and IMMUNE SYSTEM. Thus, only using the ignorance statements themselves, we have already found a set of articles and sentences for the researchers to review for a potential research topic.

**Table 9:**
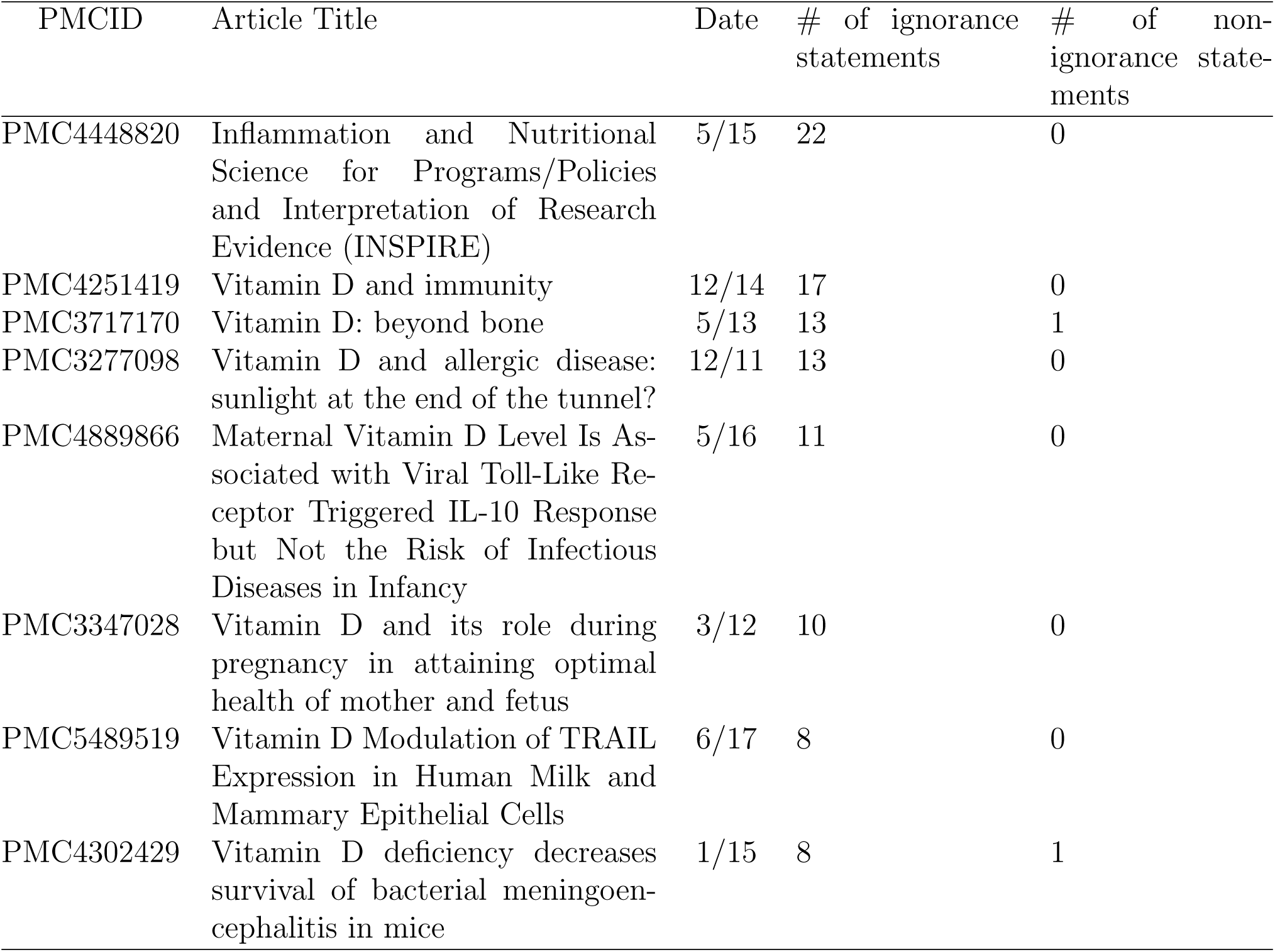
Articles with the most ignorance statements: The top eight articles for vitamin D and immune system in order of the most ignorance statements.

| PMCID | Article Title | Date | # of ignorance statements | # of non-ignorance statements |
| --- | --- | --- | --- | --- |
| PMC4448820 | Inflammation and Nutritional Science for Programs/Policies and Interpretation of Research Evidence (INSPIRE) | 5/15 | 22 | 0 |
| PMC4251419 | Vitamin D and immunity | 12/14 | 17 | 0 |
| PMC3717170 | Vitamin D: beyond bone | 5/13 | 13 | 1 |
| PMC3277098 | Vitamin D and allergic disease: sunlight at the end of the tunnel? | 12/11 | 13 | 0 |
| PMC4889866 | Maternal Vitamin D Level Is Associated with Viral Toll-Like Receptor Triggered IL-10 Response but Not the Risk of Infectious Diseases in Infancy | 5/16 | 11 | 0 |
| PMC3347028 | Vitamin D and its role during pregnancy in attaining optimal health of mother and fetus | 3/12 | 10 | 0 |
| PMC5489519 | Vitamin D Modulation of TRAIL Expression in Human Milk and Mammary Epithelial Cells | 6/17 | 8 | 0 |
| PMC4302429 | Vitamin D deficiency decreases survival of bacterial meningoen- cephalitis in mice | 1/15 | 8 | 1 |

Our ignorance taxonomy includes 13 categories of unknowns that can help researchers narrow their search. Note that the COVID-19 search engine [20] only distinguishes between two categories, scientific challenges and new directions. So we continued to narrow our search using ignorance-category enrichment, since choosing the IMMUNE SYSTEM was still quite a large topic with lots of ignorance statements. Understanding and tracing ignorance categories over time can both help the researchers narrow in on a research topic and also show how the questions in a field are asked more broadly. VITAMIN D ignorance statements in general employed a wide-range of different ignorance categories (see Figure 11), spanning all the thirteen categories of ignorance. Ten were enriched in VITAMIN D ignorance statements as compared to all ignorance statements (see the green highlights in Figure 11). For example, *unknown/novel* was enriched in this domain, pointing to large unknowns about the context of VITAMIN D in pregnancy and fetal development. To also understand how these questions changed over time, the bubble plot for IMMUNE SYSTEM and VITAMIN D ignorance statements (see Figure 12) showed that *unknown/novel* was spread out amongst the different articles. This suggests that researchers have not resolved their broadest unknowns in this field over time, in which case it may be a good knowledge goal area for a research topic. Thus the researchers can continue to narrow in on a research topic not only with a biomedical concept, such as IMMUNE SYSTEM, but also with an ignorance category, such as *unknown/novel*.

**Figure 11:**
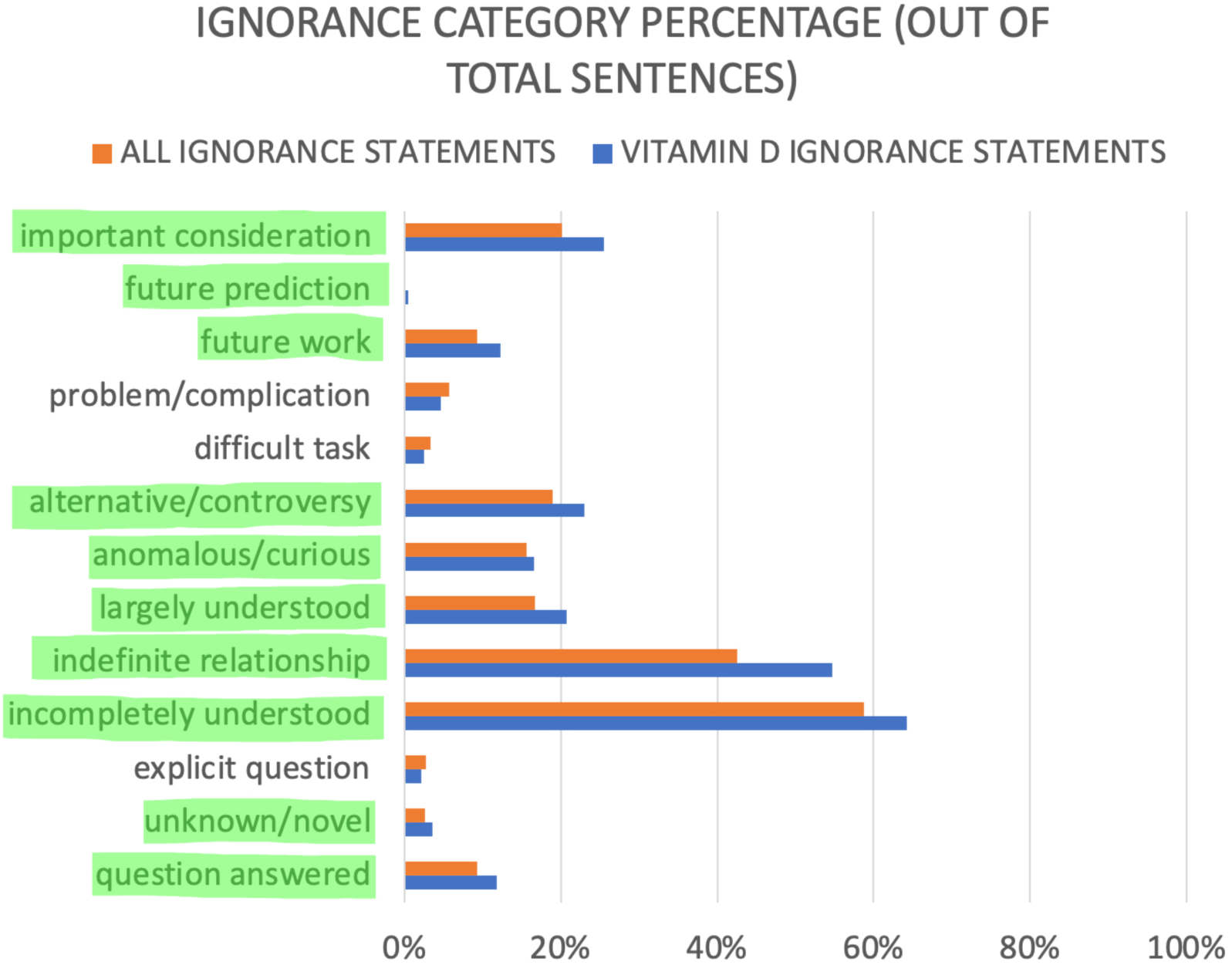
Ignorance-category enrichment: Ignorance vitamin D sentences compared to all ignorance sentences. The 10 categories highlighted in green were enriched.

**Figure 12:**
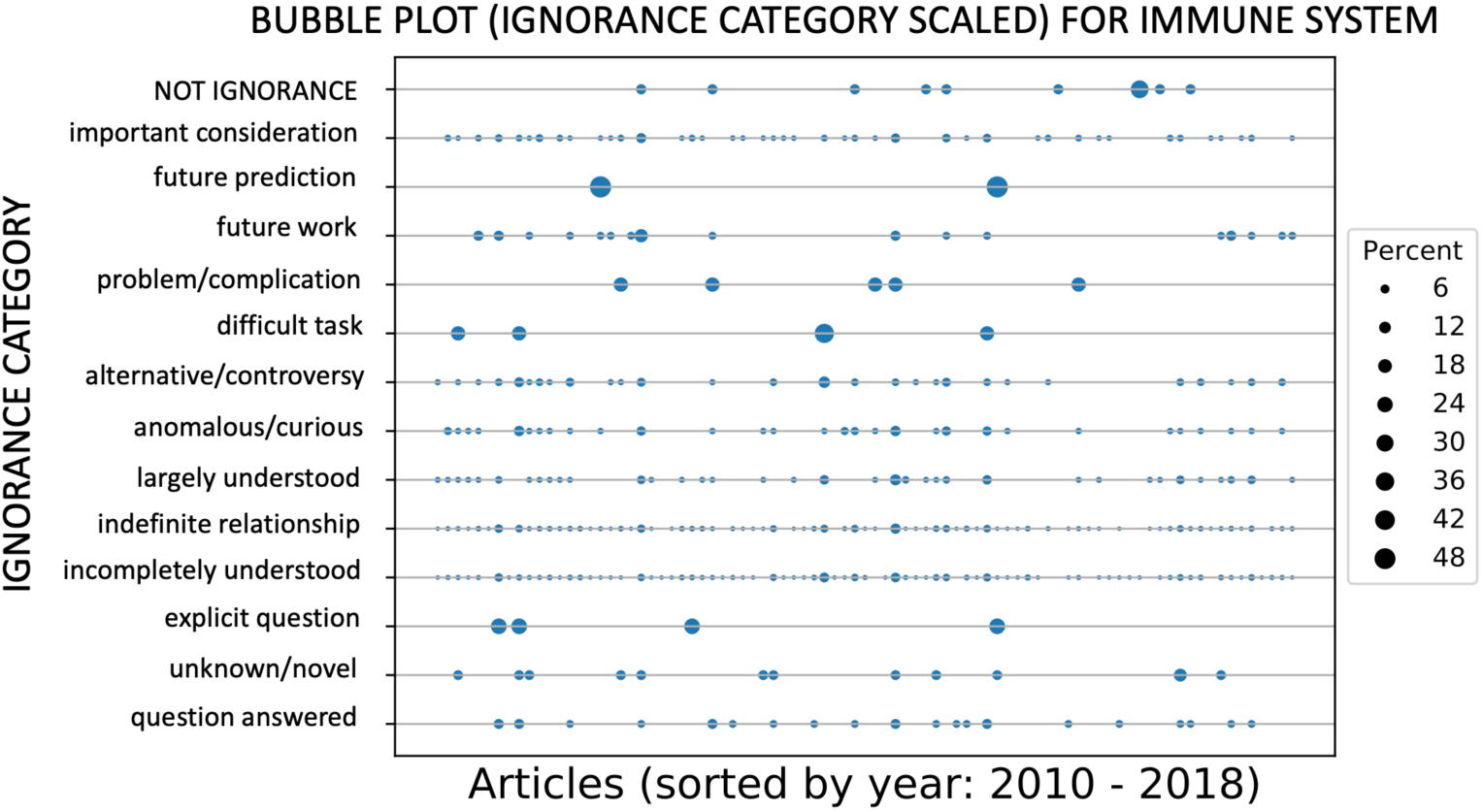
How ignorance changes over time: A bubble plot of vitamin D and immune system sentences (including non-ignorance sentences). The x-axis is the articles sorted by time. The y-axis is the ignorance categories. Each bubble represents the portion of sentences in each article in that ignorance category (scaled by the amount of total ignorance sentences in the category). For example, *future prediction* only appears in two different articles and is basically split in half between both.

We chose to dive deeper into the *unknown/novel* category for VITAMIN D and IMMUNE SYSTEM, with the goal to find pertinent questions as an example of exploration by topic. There were many *unknown/novel* ignorance sentences to investigate here. Below are some ignorance sentences (lowercase) from this set with the biomedical concepts capitalized and the ignorance lexical cues underlined:

1. “in the last five years, there has been an explosion of published data concerning the IMMUNE <u>effects</u> of VITAMIN D, yet little is known in this regard about the specific IMMUNE <u>effects</u> of VITAMIN D during PREGNANCY.” (PMC3347028)
2. “t<u>hese</u> results <u>describe</u> <u>novel</u> mechanisms and new concepts with regard to VITAMIN D and the IMMUNE SYSTEM and suggest therapeutic targets for the CONTROL of AUTOIMMUNE diseases.” (PMC3717170)
3. “however, findings regarding the combined e<u>ffec</u>t<u>s</u> of PRENATAL and POSTNATAL VITAMIN D status on fs [food sensitization], two of the most <u>critical</u> periods for IMMUNE SYSTEM DEVELOPMENT (19,20), are <u>unclear</u>.” (PMC3773018)
4. “it has an important <u>role</u> in BONE HOMEOSTASIS, BRAIN DEVELOPMENT and MODULATION OF the IMMUNE SYSTEM and yet the impact of ANTENATAL VITAMIN D deficiency on infant outcomes is poorly understood.” (PMC4072587)
5. “background: VITAMIN D is known to <u>affect</u> IMMUNE function; <u>however</u> it is <u>uncertain</u> <u>if</u> VITAMIN D <u>can</u> alter the IMMUNE RESPONSE towards the persistent herpesviruses, EBV and CMV.” (PMC4113768)

(Note that not all biomedical concepts were recognized by the biomedical concept classifiers.) The overall research topic or knowledge goal based on these statements was the need to explore the relationship between VITAMIN D and the IMMUNE SYSTEM especially in pregnancy. The same methods can be used for the other top enriched concepts including BRAIN DEVELOPMENT (data not shown). For BRAIN DEVELOPMENT, the overarching knowledge goal was the need to determine if VITAMIN D and BRAIN DEVELOPMENT were truly linked. Thus, from querying the ignorance-base for the topic VITAMIN D, the researchers now have knowledge goals to pursue in specific concept areas. (see Figure 8). Our exploration by topic methods provided multiple starting points for this research, in more depth than the standard approach and the COVID-19 search engine [20] can supply.

### 4.3. Connecting experimental results (e.g., a gene list) to ignorance statements can identify questions that may bear on it, providing new avenues for exploration, potentially from other fields

Similar to exploration by topic, exploration by experimental results provided the ignorance context for a gene list as possible future work for the researchers. Note that this was made possible by the OBOs and that neither the standard literature approach nor the COVID-19 search engine [20] have this capability. Connecting a vitamin D and sPTB gene list from a paper [36] to ignorance statements found a new avenue for exploration, BRAIN DEVELOPMENT, that was not mentioned in the paper, and pointed to an implied field, neuroscience, as a possible source for answers.

Following the exploration by experimental results pipeline (see Figure 4), the 43 genes mapped to 782 OBO concepts. These OBOs connected to 51,868 sentences (1,590 articles) that included 17,586 unique OBO concepts (88% of the total unique OBO concepts), of which, 33,885 sentences (1,537 articles) were ignorance statements with 11,711 unique OBO concepts (59% of the total unique OBO concepts). This suggests that the majority of sentences connected to these genes were ignorance statements (65%). These data can be explored by topic using the OBO list, but we focused on the three new analyses. The three new analyses were helpful to digest both the many OBO concepts and the many statements of ignorance connected to the gene list to provide areas of research to explore in future work.

With the gene list connected to so many concepts (782), combining gene list coverage and ignorance enrichment helped prioritize concepts to explore (see Table 10). The highest covered OBO concept was PROTEIN CODING GENE (SO:0001217). In the top 25 most covered OBO concepts, all concepts were established information, currently studied, or not significant (see Figure 3). (None were emerging topics.) Note that some had no information from the literature-side, meaning no conclusions could be drawn based on the current information. In fact, all 18 concepts enriched in ignorance, were also enriched in all gene list sentences, including: PROTEIN CODING GENE, INNATE IMMUNE RESPONSE, GENE, IMMUNE RESPONSE, and BRAIN in the top 25. As before, concepts related to IMMUNE SYSTEM and BRAIN surfaced after ignorance enrichment. These concepts were ripe for exploration in relation to the gene list to find knowledge goals that may bear on them.

**Table 10:** Gene list coverage enrichment information for the top 25: NO INFO stands for NO INFORMATION meaning that the ontology term exists in PheKnowLator and is connected to our gene list, but there were no sentences that contained it on the literature side. *Statistically significant with FDR but not family-wise error.

| OBO ID | OBO label | Gene Coverage | Enriched in all gene list sentences | Enriched in Ignorance |
| --- | --- | --- | --- | --- |
| SO:0001217 | protein coding gene | 37 | YES* | YES |
| GO:0005515 | response to virus | 23 | NO INFO | NO INFO |
| CL:0000094 | granulocyte | 19 | YES | NO |
| GO:0005886 | plasma membrane | 18 | YES | NO |
| UBERON:0000178 | blood | 17 | YES | NO |
| UBERON:0002371 | bone marrow | 15 | YES | NO |
| CL:0000576 | monocyte | 14 | YES | NO |
| GO:0005576 | extracellular region | 14 | YES | NO |
| GO:0005615 | extracellular space | 13 | YES | NO |
| CL:0000775 | neutrophil | 13 | YES | NO |
| CHEBI:2504 | aflatoxin B1 | 12 | YES | NO |
| CHEBI:39867 | kidney | 11 | NO INFO | NO INFO |
| GO:0045087 | innate immune response | 11 | YES | YES* |
| GO:0070062 | extracellular exosome | 11 | YES | NO |
| GO:0016021 | bone marrow | 10 | NO INFO | NO INFO |
| SO:0000704 | gene | 9 | YES | YES |
| GO:0005829 | cytosol | 9 | YES | NO |
| CL:1001608 | foreskin fibroblast | 7 | NO | NO |
| UBERON:0001332 | prepuce of penis | 7 | YES* | NO |
| GO:0005737 | cytoplasm | 7 | YES | NO |
| GO:0006955 | immune response | 7 | YES | YES |
| CL:0000765 | erythroblast | 7 | NO | NO |
| UBERON:0000955 | brain | 6 | YES | YES |
| GO:0035580 | lead(0) | 6 | NO INFO | NO INFO |
| CL:0000771 | eosinophil | 6 | YES | NO |

Combining other canonical enrichment methods with ignorance enrichment also helped prioritize the many OBO concepts produced by our gene list (see Figure 13 focusing only on the gene ontology). DAVID [144, 145] found 42 of the 43 genes and mapped them to 159 GO concepts. 51 of those were enriched and 30 were contained in sentences found in the ignorance-base. Of those 30, 19 were contained in gene list statements and 11 had no information. Of those 19, 17 had at least one ignorance statement, and the concepts were mainly related to the immune system. (Two concepts, RESPONSE TO STRESS (GO:0006950) and MULTI-ORGANISM PROCESS (GO:0051704) had no ignorance statements.) The ignorance statements for the 17 concepts can be explored to provide more information to the canonical enrichment methods and their respective knowledge-bases.

**Figure 13:**
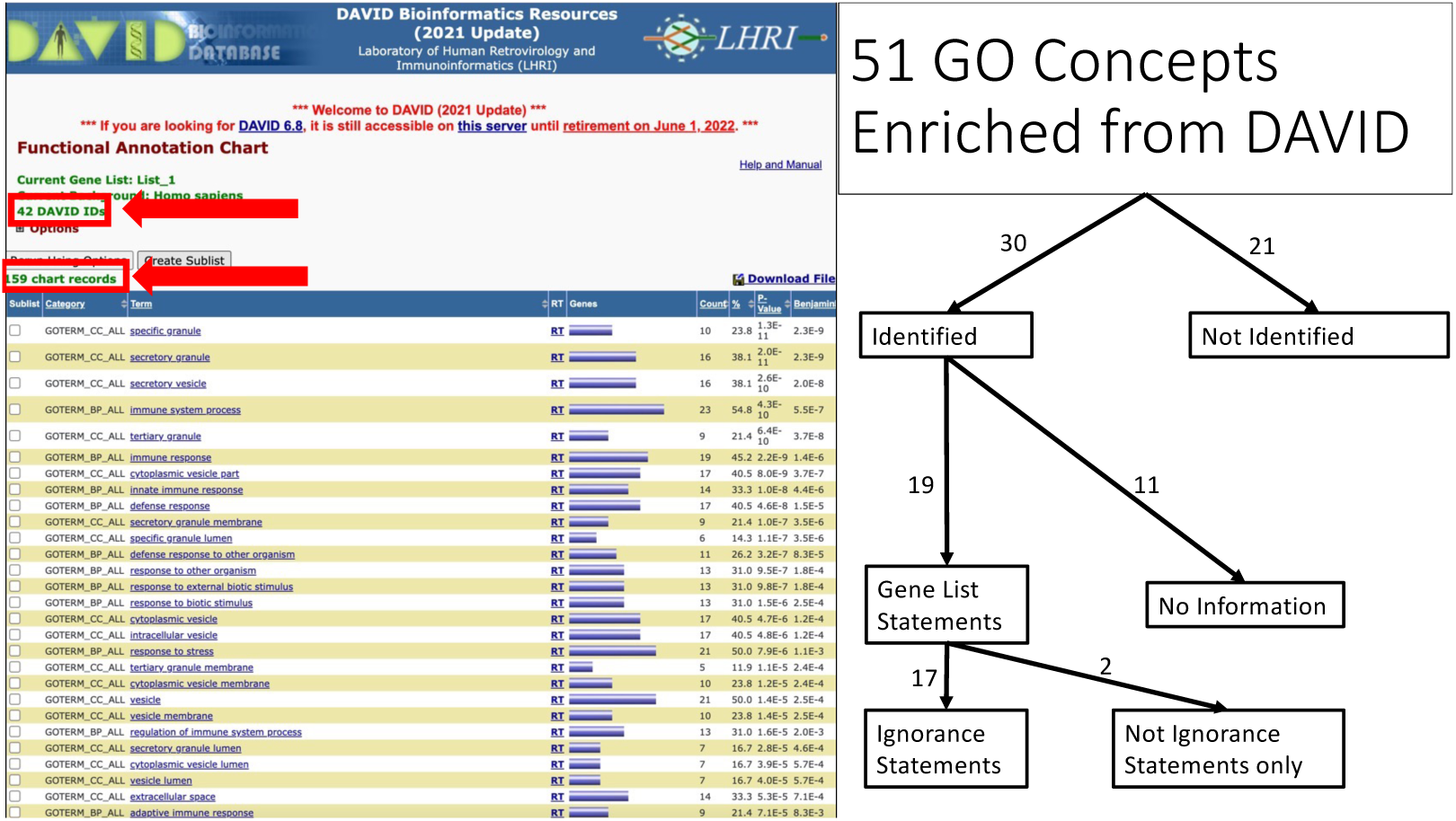
Enhancing canonical enrichment analysis using the ignorance-base: DAVID enrichment analysis for the gene ontology (GO) in relation to the ignorance-base. The DAVID initial analysis is on the left with 42 of the 43 genes found in DAVID mapping to 159 GO concepts. The right is a breakdown of where the 51 enriched GO concepts from DAVID fall within the ignorance-base.

To find out whether our ignorance lens could augment canonical methods, we compared our ignorance approach to the canonical approach. When comparing ignorance enrichment in GO to DAVID, we found more ignorance in general compared to established information. 3,173 GO concepts were enriched in ignorance with 159 in DAVID (see Figure 14). Intriguingly, the overlap between the two analyses was small: 60. If we look at enrichment it was even smaller: only two concepts overlapped. Potentially this makes sense as we were enriching for the opposite things: ignorance vs. established knowledge. On the knowledge-side, most enriched concepts from DAVID pertained to the immune system, which was the main focus of Yadama *et al.*, [36]. The ignorance-side found more general biological processes, and two concepts from the overlap, IMMUNE RESPONSE and INNATE IMMUNE RESPONSE, also pertained to immunity. These concepts were currently studied.

**Figure 14:**
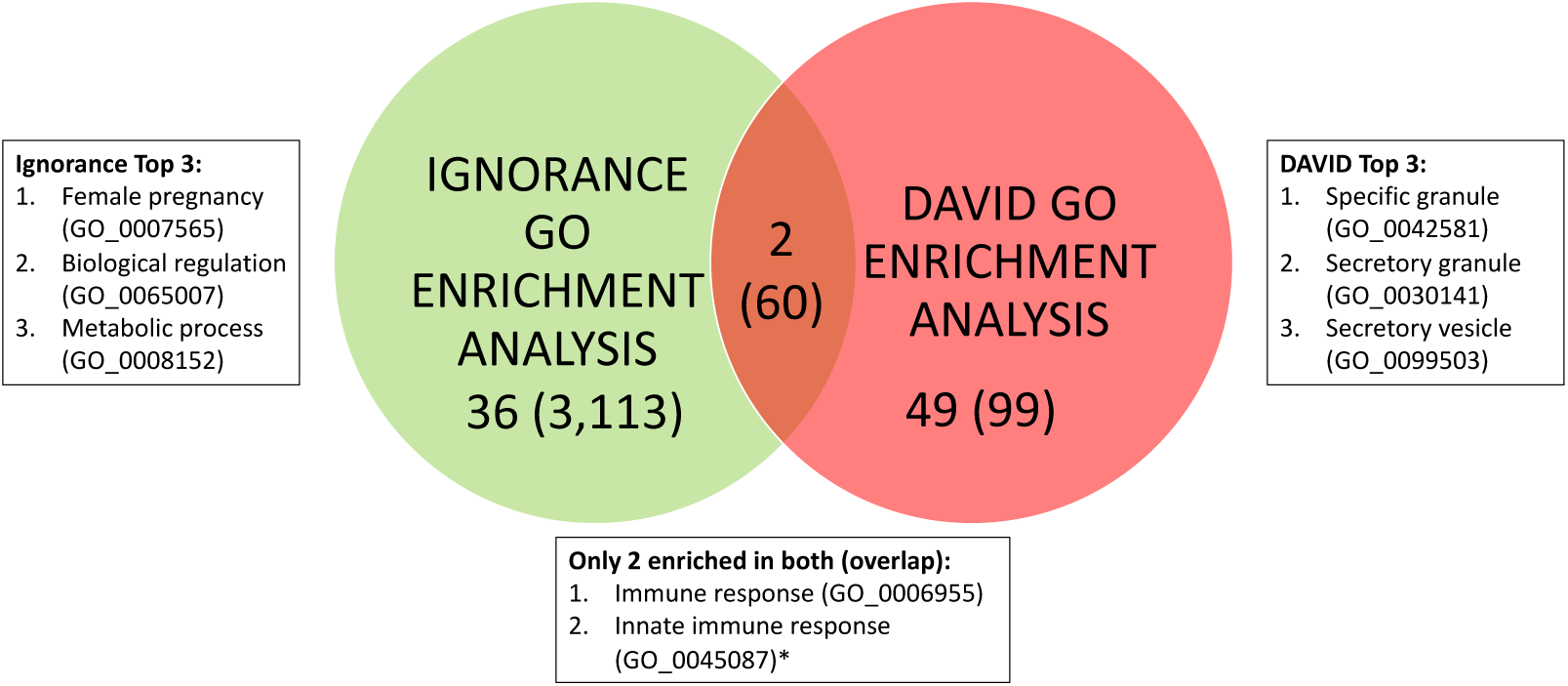
Comparison of DAVID and ignorance enrichment: A Venn diagram of gene ontology enrichment between DAVID (pink) and the ignorance-base (green). In parentheses are the total number of concepts found in each category without enrichment. Next to each bubble are the top three concepts for each enrichment method. The concepts in the middle are the overlap. *Statistically significant with FDR but not family-wise error.

Looking at the ignorance enriched concepts only, we achieved our goal of finding a new avenue to investigate that was not mentioned by the paper [36], namely the brain. Further, it also provided a different field to examine for answers, namely neuroscience. The top three ignorance-enriched GO concepts included FEMALE PREGNANCY, BIOLOGICAL REGULATION, and METABOLIC PROCESS (see Figure 13). Broadening this exploration beyond GO, there were 130 total ignorance enriched concepts for this gene list, including the 38 from GO. There were still some immune related concepts including (in order of enrichment): IMMUNE SYSTEM, IMMUNE RESPONSE, SKELETON OF MANUS (autoimmune), INTERLEUKIN-1 FAMILY MEMBER 7*, and INNATE IMMUNE RESPONSE*. (The * means that they were statistically significant with FDR but not familywise error.) As mentioned above, there was still future work to understand the IMMUNE SYSTEM in relation to sPTB and VITAMIN D [36], and the ignorance approach provided specific ignorance statements to explore it. Even more striking though was the number of ignorance enriched concepts related to the BRAIN (12): BRAIN, BRAIN DEVELOPMENT, COGNITION, NERVOUS SYSTEM DEVELOPMENT, NEURON, LEARNING, SYNAPSE, NERVOUS SYSTEM, CENTRAL NERVOUS SYSTEM, NEUROTRANSMITTER*, NEURAL TUBE*, and NEUROGENESIS. Neither Yadama *et al.*, [36] nor DAVID [144, 145] derived any brain-related concepts from this gene list, signifying that this association was not yet established knowledge.

The developing brain may be an emergent topic related to vitamin D and spontaneous preterm birth, and neuroscience may shed light on these connections. Along these lines, we present some excerpts (lowercase) from five papers to explore below (the biomedical concepts are capitalized and the ignorance lexical cues are underlined):

1. “this AREA OF the BRAIN, specifically the FRONTAL CORTEX, is important for LANGUAGE, MEMORY and higher order COGNITIVE functioning, including purposeful, goal-directed behaviours which are often referred to as executive functions.^4^ the <u>importance</u> of adequate dha [docosahexaenoic acid] during this key period of BRAIN DEVELOPMENT is <u>indicated</u> in studies of preterm infants who are denied the full GESTATION period to accumulate DHA” (PMC4874207)
2. “discussion: we review r<u>elevant</u> literature suggesting in utero inflammation <u>can</u> <u>lead</u> to PRETERM labor, <u>while</u> <u>insufficient</u> development of the GUT-BLOOD–BRAIN barriers <u>could</u> permit exposure to potential neurotoxins.” (PMC3496584)
3. “a major Intake of DHA in the BRAIN happens in the last TRIMESTER of PREGNANCY; <u>therefore</u>, preterm infants are disadvantaged and have decreased BRAIN concentration of this vital lcpufa [long-chain polyunsaturated fatty acid].” (PMC3607807)
4. “at present, preterm infants have a limit of viability (50% survival rate) of around 23–24 weeks ga [gestational age] so post-NATAL nutrition will always be introduced during the second major phase of BRAIN growth, resulting in differences mainly in WHITE MATTER.” (PMC3734354)
5. “the most likely explanation <u>seems</u> to be <u>related</u> to the timing of the nutrition event, <u>since</u> the infants were BORN at term <u>rather</u> than preterm when different developmental processes are occurring in the BRAIN.” (PMC3734354)
6. “the period between their PRETERM BIRTH and term BIRTH at 40 weeks, a time when the major BRAIN spurt is occurring), was spent ex utero in these infants; this exposure to environmental <u>influences</u>, at an early stage of BRAIN DEVELOPMENT, might be expected to increase their vulnerability to dietary <u>effects</u>.” (PMC3734354)
7. “the authors conclude by saying that reducing the energy deficit by improving early nutrition in preterms may improve the growth and maturation of the BRAIN.” (PMC3734354)
8. “iron status, more commonly assessed in PREGNANCY, is not only important in HEMATOPOESIS and NEUROLOGICAL and COGNITIVE DEVELOPMENT^9^ but plays a <u>crucial</u> <u>role</u> in CARNITINE SYNTHESIS,^10^ <u>although</u> CARNITINE precursors <u>may be</u> more <u>important</u>.^11^ zinc is an important COFACTOR for more than 300 <u>identified</u> zinc metalloenzymes.^12^ zinc insufficiency in late PREGNANCY disrupts NEURONAL REPLICATION and SYNAPTOGENSIS,^13^ and maternal deficiency is <u>associated</u> with decreased dna, rna, and protein content of the f1 BRAIN.^14^ zinc deficiency <u>affects</u> one in five world inhabitants.^14^ZINC supplementation reduces the <u>risk</u> of PRETERM BIRTH, though not sga [small for gestational age].^14^ VITAMIN D deficiency is under investigation for its <u>role</u> in protection against dm [diabetes mellitus], cv [cardiovascular], some ca [cancers], osteoporosis, and optimization of IMMUNE function.^15^ VITAMIN D might be an important mediator in GUT HOMEOSTASIS and in signaling between microbiota and host.^16^ the INTESTINAL microbiome in both newborns and LACTATING mothers <u>influences</u> infant and childhood FOOD allergy and eczema.” (PMC4268639)

(Note that not all biomedical concepts were recognized by the biomedical concept classifiers. Also, the numbers in the sentences represent citations, which were superscript in the original article but were flattened for processing.)

In the paper, Yadama *et al.*, [36] focused on the mother’s immune system, but it is possible that this brain connection is instead focused on the effects of the spontaneous preterm birth on the offspring. Thus, a potential knowledge goal for the authors based on our analysis was to explore the connections between maternal VITAMIN D levels and spontaneous preterm birth through the maternal IMMUNE SYSTEM and the effects on the BRAIN DEVELOPMENT of the offspring. Exploring these connections would greatly impact mothers and their offspring globally. The ignorance-base provided a novel avenue (and field), BRAIN DEVELOPMENT (neuroscience), along with specific knowledge goal statements, that the authors can explore in future work based on their initial gene list. Our exploration by experimental results method contextualized experimental results in the ignorance landscape, providing multiple avenues for future research, the immune system and the brain.

## 5. Discussion

Focusing on ignorance statements through our ignorance-base and exploration methods led to new research avenues that could help accelerate research. Further, the ignorance-base is more than just a literature search engine similar to Lahav *et al.*, [20]; it also provided insights, summaries, and visualizations based on topics and experimental results. Its focus on knowledge goals and its grounding in the OBOs has helped our ignorance-base to find areas of research with many questions and to identify fields of study that may contain answers. The ignorance-base predicted areas of research that were currently studied and an emerging topic with a corresponding field that may prove fruitful to help find answers.

The exploration by topic method showed that vitamin D may play an important role in the immune system, respiratory system, and brain development (see Figures 8-12 and Table 9). Corroborating these findings after 2018, when our corpus ended, recent review articles [10, 11, 12, 13, 14] included these areas as future work. These review articles required many hours of reading and synthesizing the literature article by article, whereas the ignorance method automatically offered not only articles, but also specific sentences that discuss knowledge goals for future work. For example, the sentence “it has an important <u>role</u> in BONE HOMEOSTASIS, BRAIN DEVELOPMENT and MODULATION OF the IMMUNE SYSTEM and yet the impact of ANTENATAL VITAMIN D deficiency on infant outcomes is poorly understood” (PMC4072587) [167] showed that further research on the impact of vitamin D on infant outcomes was needed. The context of the sentence is also important: it comes from the abstract objective section of a 2014 study in Rural Vietnam. Because our ignorance approach allows sorting of statements by time and section, we showed that since 2014, more research has been conducted on this topic [12]. Further, we can track how research questions emerge using our ignorance taxonomy (see Figure 12). Even this smaller-scale effort, limited to one broad topic and the years 1939-2018, demonstrated that we can map the landscape of our collective scientific ignorance and track how research questions evolve over time. Ideally, future work would create an ignorance-base over the entire body of scientific literature to provide this resource to researchers, students, funders, and publishers.

We further demonstrated that the ignorance-base and exploration by experimental results method can find an emerging topic (see Figures 4, 13, 14, and Table 10). Ignorance enrichment of the 43 genes in common between vitamin D and spontaneous preterm birth (sPTB) [36] found many concepts that relate to the brain and some that relate to the immune system (as found by [36]). This suggested that brain development could be an emerging topic in relation to vitamin D and sPTB. For example, consider the ignorance statement: “discussion: we review <u>relevant</u> literature suggesting in utero inflammation <u>can</u> <u>lead</u> to PRETERM labor, <u>while</u> <u>insufficient</u> development of the GUT-BLOOD–BRAIN barriers <u>could</u> permit exposure to potential neurotoxins” (PMC3496584) [168]. This sentence ties all the relevant concepts together by suggesting that “vitamin D may be causing in utero inflammation leading to preterm labor; due to the preterm labor the gut-blood-brain barrier may develop incompletely, which in turn exposes the fetus to potential neurotoxins”. Although this article was not cited by *et al.*, [36], it may posit a new knowledge area that needs to be explored further. Lastly, researchers could look to the field of neuroscience to help find relevant information to some of these knowledge goals. In consultation with our prenatal nutrition expert, here are some potential questions that could be explored:

1. What is the association between development of the gut-blood-brain barrier and whole-body inflammation and neuroinflammation in the context of fetal development?
2. How do the 43 genes relate to offspring brain development? Are any of them specifically related to offspring brain function?
3. What are the effects of vitamin D on lifecourse brain development generally?
4. How does spontaneous preterm birth effect offspring brain development compared to those born at term? Are there any remedies for said effect? Does nutrition play a role?
5. How does the gestational timing of nutrition and supplement exposure affect offspring brain development? What role do iron and zinc play in brain development?

To corroborate our findings, looking at the literature further showed that vitamin D and brain development may in fact be an emerging topic since 2018, the last year of our data. The connection between vitamin D and brain development has only recently been studied extensively. Looking for recent papers on “vitamin D”, “brain development”, and “spontaneous preterm birth” in the literature (Google Scholar search on 9/13/2022), many review articles appeared ([10, 11, 12, 13, 14]) that discuss the impact of vitamin D on maternal and fetal health (see Figure 3 in [10] and our adapted Figure 1). All of these studies drew links between vitamin D, the immune system, and sPTB, and acknowledge at least one link between vitamin D and brain development. A 2022 review article stated that “recently, extensive scientific literature has been published determining the role of vitamin D in brain development” [10]. Note that these articles contain mentions of controversies and other types of ignorance statements, which also point to other areas of investigation. There appears to be room for exploration around the connections between vitamin D, sPTB, and brain development. Our ignorance approach could help automate review articles in finding the emerging topics to study. Understanding the ignorance-context around a set of genes in combination with the knowledge-context can help push the boundaries of our current understanding.

In general, we showed that ignorance-bases and knowledge-bases can enhance and complement each other. The ignorance-base itself was built upon a knowledge-base, PheKnowLator [26, 27]. We also utilized DAVID [145] as a comparison knowledge-base (see Figures 13 and 14) as well as other canonical methods (gene list coverage) to help prioritize the most relevant biomedical concepts (see Table 10). These analyses were made possible by grounding our ignorance-base in the OBOs, which allowed us to connect our ignorance-base to other knowledge-bases. At the same time, we did not use any of these methods to their fullest potential. First, only six ontologies overlapped between the biomedical classifiers and PheKnowLator, which limited the expansion of relevant concepts. Second, the method to create the OBO concept lists from both the vitamin D topic and the gene list were not very sophisticated (using only one step via the relations ontology). Finally, we did not use the knowledge-bases to determine if any ignorance statements have been answered. This is quite a hard problem that Lahav *et al.*, [20] also did not tackle. All of these limitations could be addressed in future work. Further, there are many other knowledge-bases and methods that can be explored in relation to the ignorance-base.

The goal of this work was to demonstrate feasibility of the ignorance methods and we recognize that more improvements can be made. Our ignorance-base was created from automatic classifiers run over 1,643 articles in the prenatal nutrition literature. Any automation of this kind adds errors and all classification tasks can be improved upon to minimize it. For the ignorance classifiers, other parameter tunings and other algorithms, such as PubMedBERT [169], may yield improved results. Lahav *et al.*, [20] used PubMedBERT along with other algorithms. However, overall performance was lower than ours. Our biomedical concept classifiers were developed using CRAFT [158, 157], a corpus of mouse articles, not prenatal nutrition, and it only included ten ontologies. Applying biomedical classifiers with more similar training data and more ontologies (*e.g.*, MONDO disease ontology and the phenotype ontology) would be beneficial (*e.g.*, PubTator [170]), although all of them have their pros and cons. We ran these classifiers over only 1,643 prenatal nutrition articles. The scale of the ignorance-base was small; ideally we would create an ignorance-base that included all articles (or at least PMCOA to start). We focused only on the prenatal nutrition literature, and future work will determine if the ignorance taxonomy and methods generalize outside of it. But the extent of overlap between our cue list and similar prior work (see Table 8) implies that our ignorance-base may translate to other biomedical domains. In terms of exploration methods, concept enrichment provided more fruitful concepts (see Figures 10 and 11) than concept frequency (see Figure 9). Another avenue to explore would be co-occurrence terms. The creation of a tool (similar to [20]) incorporating more data analyses and visualizations techniques into a user-interface that allows researchers to interact with the system could make the ignorance-base easier to adapt to new environments. Future work could combine these efforts. Even with all these limitations, the current ignorance-base showed its power to find new research avenues to explore, providing insights, summaries, and visualizations beyond prior work [20]. We have just barely scratched the surface of what it can do. Collaborating with experts on vitamin D, delving into the topics introduced here, creating new methods, exploring other topics, and contextualizing other experimental results are obvious extensions of this work.

There is also future work in relation to the ignorance corpus and classifiers. Highlighting the lexical cues for the annotators before annotation could have biased the annotators. We did not measure this effect because our annotators found the task infeasible when the lexical cues were not highlighted. Even still, with the highlights, the annotators continued to find new lexical cues, which further extended the reach of the classifiers. We conducted an ablation study that determined the importance of the lexical cues as features for classifying ignorance statements (see Table 7). As mentioned, our annotators found them helpful. Lahav *et al.*, [20] agreed that the task was quite difficult with misleading keywords and so they added in sentences without any of them. In contrast, we found that a larger cue list (2,513) was more resilient to error and helped our classifiers discover many more lexical cues (added in 5,637). Our IAAs were comparable to [20], in the 80% range. More data can always be annotated both to improve the current annotations and to add data from other fields besides prenatal nutrition. Confirming the ignorance taxonomy and classifiers generalize beyond prenatal nutrition will allow for the creation of a larger ignorance-base. More work needs to be done, but we showed that our lexical cues overlapped with prior work in other domains (see Table 8, hinting at the generalizability beyond our work here.

We demonstrated that a focus on ignorance statements through our ignorance-base and exploration methods can lead students, researchers, funders, and publishers to research avenues that are currently being studied or are emerging topics. Research begins from a foundation of established knowledge, and many knowledge-bases and ontologies exist to provide that. However, research continues through a process of posing questions and creating hypotheses to analyze and explore what is not yet understood. To facilitate that, we present the first ignorance-base based on knowledge goals and OBOs, along with two new exploration methods that provided insights, summaries, and visualizations of statements of unknowns, controversies, and difficulties needing resolution in future work. Just as the literature contains both knowledge and ignorance, so too can both knowledge-bases and ignorance-bases help researchers navigate the literature to find the next important questions or knowledge gaps.

## 6. Conclusion

Our ultimate goal was to create an ignorance-base and exploration methods to enable students, researchers, funders, and publishers to find the next important scientific questions or knowledge gaps. By augmenting and streamlining the manual work of literature reviews, we can help direct research to focus on important quesitons and possible answers. The exploration by topic method not only found new avenues for exploration for researchers interested in vitamin D using our novel method of ignorance enrichment (the immune system, respiratory system, and brain development), but also elucidated how questions were asked and how that changed over time using our novel method of ignorance-category enrichment. Our exploration by experimental results method found an emerging topic (brain development) with specific knowledge goal statements to pursue that bear on a sPTB and vitamin D gene list. Further, the findings suggested a field (neuroscience) to look to for answers. These questions (and subsequent answers) have high potential to positively impact the health of pregnant women and their offspring globally. The importance of questions and knowledge goals in research is well established, and our ignorance-base and exploration methods bring these to the forefront to help researchers explore a topic and experimental results in the context of our collective scientific ignorance. The scientific endeavor rests on our continuous ability to ask questions and push research farther as we learn more knowledge. To paraphrase Confucius,“Real knowledge is to know the extent of one’s ignorance” (Analects 2:17). In the right context, ignorance is a source of wisdom.

## Declaration

## Acknowledgements

We would like to acknowledge the BioFrontiers Computing Core for computing resources and support, especially Jonathon Demasi. The authors would like to thank Harrison Pielke-Lombardo for his updates to the Knowtator tool; William A. Baumgartner Jr. for his help with the literature and taxonomy work; and Tiffany J. Callahan for many discussions about this work.

## Availability of data and materials

Code for the ignorance-base and exploration methods can be found at: https://github.com/UCDenver-ccp/Ignorance-Base. The expanded ignorance corpus can be found at: https://github.com/UCDenver-ccp/Ignorance-Question-Corpus with all associated code and models at: https://github.com/UCDenver-ccp/Ignorance-Question-Work-Full-Corpus. Code for concept recognition of the OBOs can be found at: https://github.com/UCDenver-ccp/Concept-Recognition-as-Translation. Our previous related work that we built upon for this work can be found at: https://github.com/UCDenver-ccp/Ignorance-Question-Work.

## Ethics approval and consent to participate

Not applicable.

## Authors’ contributions

Mayla **R. Boguslav:** Conceptualization, Methodology, Data Curation, Software, Validation, Writing - Original Draft. **Nourah M. Salem:** Methodology, Software, Validation, Writing - Review Editing. **Elizabeth K. White:** Data Curation, Validation, Writing - Review Editing. **Katherine J. Sullivan:** Data Curation, Validation, Writing - Review Editing. **Stephanie P. Araki:** Data Curation, Validation. **Michael Bada:** Data Curation, Writing - Review Editing. **Teri L. Hernandez:** Supervision, Writing - Review Editing. **Sonia M. Leach:** Supervision, Writing - Review Editing. **Lawrence E. Hunter:** Supervision, Writing - Review Editing.

## Appendix A. Supplementary File on Corpus Information

Details on the corpus specifically.

In annotating, K.J.S. and M.R.B. decided that one article (PMC4869271) was too difficult to annotate for this task because:

- The audience seemed to be public health workers whereas this task was more focused on researchers via scientific articles. This may be a true implementation paper.
- The article was full of quotes, making it very difficult to determine what was ignorance. Many of the quotes were also informal, meaning that it was unclear what was a quote or not. This further made the article difficult to follow.
- The article was most likely more of a technical report, public health education, government, and/or an interview rather than a scientific article based on the publication types from the NIH [171]. There did not appear to be much basic research in it.
- It was rather difficult to determine what was ignorance in the study itself versus the implementation of the policies being presented. The majority of the article seemed factual because it was mainly quotes from the public health workers.
- Neither K.J.S. nor M.R.B. were confident in their annotations of the article.

Thus, this article was excluded from the final corpus.

**Figure A.15:**
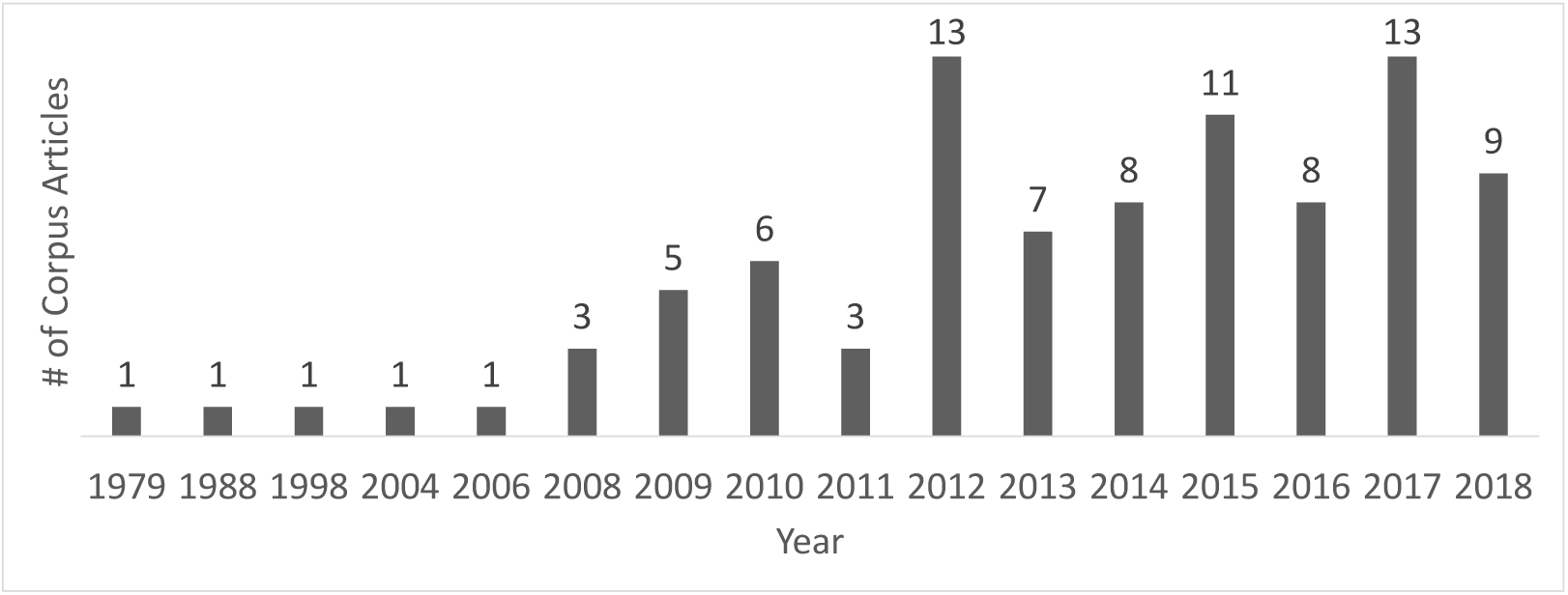
Article date distribution for the ignorance corpus (1979-2018).

**Table A.11:** Annotation Statistics Per Ignorance Category: Total number of lexical cue annotations in all articles and statistics per ignorance category. SCOPE is the number of sentences that contain at least one ignorance lexical cue. Note that all categories except for ALL CATEGORIES and SCOPE (have 1) have zero minimum number of annotations. Note max = maximum.

| Category | # total annotations | average # annotations per article | median # annotations per article | max # annotations |
| --- | --- | --- | --- | --- |
| answered question | 619 | 6.8 | 7 | 23 |
| unknown/novel | 328 | 3.6 | 2 | 20 |
| explicit inquiry | 177 | 1.95 | 1 | 27 |
| incompletely understood | 6471 | 71.11 | 67 | 329 |
| indefinite relationship | 3908 | 42.95 | 36 | 165 |
| largely understood | 1695 | 18.63 | 15 | 108 |
| anomalous/curious | 1012 | 11.12 | 9 | 39 |
| alternative/controversy | 1519 | 16.69 | 12 | 94 |
| difficult task | 320 | 3.52 | 2 | 25 |
| problem/complication | 532 | 5.85 | 3 | 29 |
| future work | 948 | 10.42 | 6 | 103 |
| future prediction | 366 | 4.02 | 2 | 52 |
| important consideration | 1786 | 19.63 | 9 | 199 |
| ALL CATEGORIES | 19681 | 216.27 | 200 | 1021 |
| SCOPE | 6700 | 73.63 | 69 | 262 |

**Table A.12:** Unique Annotation Statistics Per Ignorance Category: Total number of unique lexical cue annotations in all articles and statistics per ignorance category. We did not include SCOPE because the number of sentences is the same. Note that all categories except for ALL CATEGORIES (has 1) have zero minimum number of unique annotations. Note that the SCOPE is not in the unique table because we only capture the scope one time no matter how many lexical cue annotations occur within it. Note max = maximum.

| Category | #<br>total<br>unique<br>anno-<br>tations | %<br>unique<br>anno-<br>tations | avg #<br>unique<br>anno-<br>tations<br>per<br>article | median<br>#<br>unique<br>anno-<br>tations<br>per<br>article | max #<br>unique<br>annota-<br>tions |
| --- | --- | --- | --- | --- | --- |
| answered question | 49 | 8% | 4.32 | 5 | 14 |
| unknown/novel | 69 | 21% | 2.53 | 2 | 9 |
| explicit inquiry | 16 | 9% | 1.21 | 1 | 8 |
| incompletely understood | 368 | 6% | 32.66 | 35 | 83 |
| indefinite relationship | 130 | 3% | 15 | 15 | 45 |
| largely understood | 106 | 6% | 7.59 | 7 | 25 |
| anomalous/curious | 63 | 6% | 4.6 | 5 | 14 |
| alternative/controversy | 100 | 7% | 7.57 | 7 | 24 |
| difficult task | 40 | 13% | 2.3 | 2 | 11 |
| problem/complication | 41 | 8% | 2.91 | 2 | 12 |
| future work | 105 | 11% | 5.34 | 4 | 21 |
| future prediction | 14 | 4% | 1.84 | 1 | 6 |
| important consideration | 101 | 6% | 6.73 | 5 | 37 |
| ALL CATEGORIES | 1202 | 6% | 94.59 | 104 | 273 |

**Table A.13:** Annotation Counts per Section: Total number of annotations by section in all articles with section delineation and statistics per article. Note that every article contained a title and none of the titles had any ignorance annotations. Note avg = average, min = minimum, and max = maximum.

| Section | # total<br>articles | # total<br>annota-<br>tions | avg # anno-<br>tations<br>per<br>article | median # an-<br>notations per<br>article | min # an-<br>notations | max # an-<br>notations |
| --- | --- | --- | --- | --- | --- | --- |
| abstract | 71 | 123 | 1.73 | 1 | 0 | 29 |
| introduction | 86 | 2649 | 30.8 | 16 | 0 | 571 |
| methods | 61 | 2880 | 47.21 | 38 | 1 | 367 |
| results | 54 | 927 | 17.17 | 6.5 | 0 | 287 |
| discussion | 57 | 4906 | 86.07 | 55 | 3 | 330 |
| conclusion | 41 | 8196 | 199.9 | 150 | 2 | 990 |

## Appendix B. Supplementary File on Ignorance Classification

More detailed Tables and Figures on Ignorance classification at both the sentence- and word-levels.

**Figure B.16:**
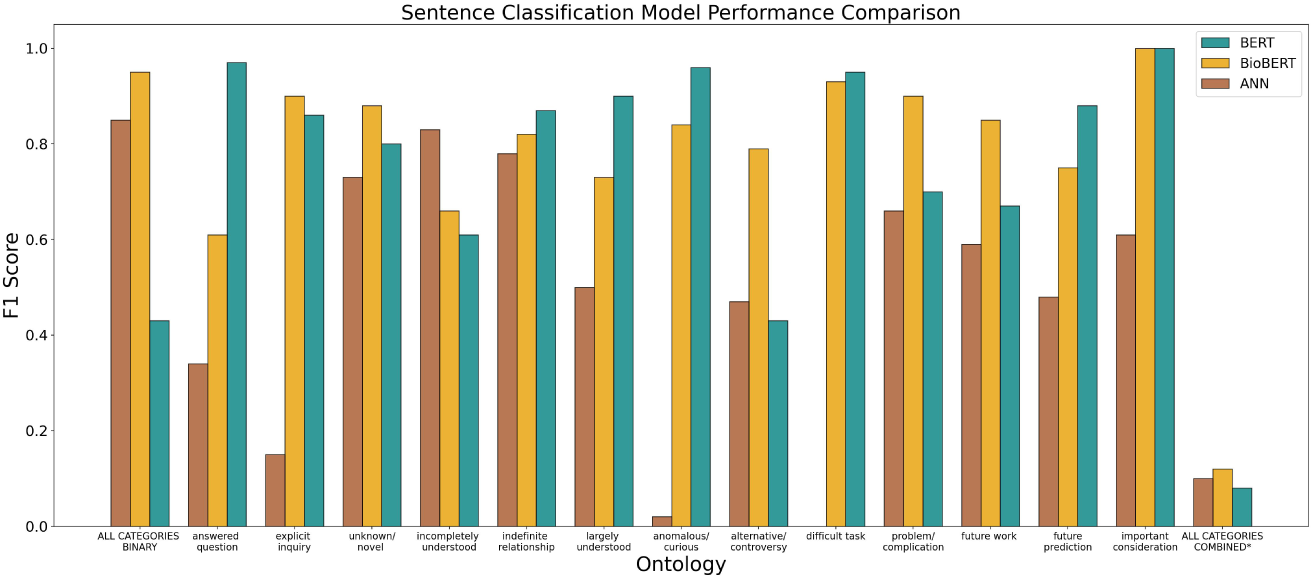
Sentence Classification summary: A bar plot summary of test F1 scores for sentence classification. *Reporting the macro-average F1 score of all the categories for one multi-classifier.

**Table B.14:** ANN sentence classification: Note that one sentence can map to more than one category and so they will not add up to the total binary. *Reporting the macro-average F1 score of all the categories for one multi-classifier.

| Ignorance Category | training<br>F1 score | training<br>support | testing F1 score | testing<br>support |
| --- | --- | --- | --- | --- |
| ALL CATEGORIES BINARY | 0.99 | 3354 | 0.85 | 2005 |
| answered question | 0.66 | 479 | 0.34 | 168 |
| explicit inquiry | 0.95 | 524 | 0.15 | 92 |
| unknown/novel | 0.87 | 347 | 0.73 | 63 |
| incompletely understood | 0.9 | 1582 | 0.83 | 225 |
| indefinite relationship | 0.86 | 1965 | 0.78 | 1072 |
| largely understood | 0.86 | 523 | 0.5 | 312 |
| anomalous/curious | 0.92 | 479 | 0.02 | 149 |
| alternative/controversy | 0.97 | 942 | 0.47 | 441 |
| difficult task | 0.88 | 342 | 0.0 | 93 |
| problem/complication | 0.68 | 324 | 0.66 | 202 |
| future work | 0.85 | 869 | 0.59 | 195 |
| future prediction | 0.93 | 178 | 0.48 | 55 |
| important consideration | 0.75 | 735 | 0.61 | 491 |
| ALL CATEGORIES COMBINED* | 0.17 | 3354 | 0.08 | 2005 |

**Table B.15:** BERT sentence classification: Note that one sentence can map to more than one category and so they will not add up to the total binary. *Reporting the macro-average F1 score of all the categories for one multi-classifier.

| Ignorance Category | training<br>F1 score | training<br>support | testing F1 score | testing<br>support |
| --- | --- | --- | --- | --- |
| ALL CATEGORIES BINARY | 0.83 | 3354 | 0.43 | 2005 |
| answered question | 0.99 | 479 | 0.97 | 168 |
| explicit inquiry | 0.87 | 524 | 0.86 | 92 |
| unknown/novel | 0.89 | 347 | 0.8 | 63 |
| incompletely understood | 0.81 | 1582 | 0.61 | 225 |
| indefinite relationship | 0.91 | 1965 | 0.87 | 1072 |
| largely understood | 0.96 | 523 | 0.9 | 312 |
| anomalous/curious | 0.93 | 479 | 0.96 | 149 |
| alternative/controversy | 0.83 | 942 | 0.43 | 441 |
| difficult task | 0.92 | 342 | 0.95 | 93 |
| problem/complication | 0.92 | 324 | 0.7 | 202 |
| future work | 0.83 | 869 | 0.67 | 195 |
| future prediction | 0.9 | 178 | 0.88 | 55 |
| important consideration | >0.99 | 735 | >0.99 | 491 |
| ALL CATEGORIES COMBINED* | 0.23 | 3354 | 0.10 | 2005 |

**Table B.16:** BioBERT sentence classification: Note that one sentence can map to more than one category and so they will not add up to the total binary.

| Ignorance Category | training<br>F1 score | training<br>support | testing F1 score | testing<br>support |
| --- | --- | --- | --- | --- |
| ALL CATEGORIES BINARY | 0.99 | 3354 | 0.95 | 2005 |
| answered question | 0.93 | 479 | 0.61 | 168 |
| explicit inquiry | 0.99 | 524 | 0.9 | 92 |
| unknown/novel | 0.88 | 347 | 0.88 | 63 |
| incompletely understood | 0.97 | 1582 | 0.66 | 225 |
| indefinite relationship | 0.88 | 1965 | 0.82 | 1072 |
| largely understood | 0.86 | 523 | 0.73 | 312 |
| anomalous/curious | 0.85 | 479 | 0.84 | 149 |
| alternative/controversy | 0.92 | 942 | 0.79 | 441 |
| difficult task | 0.91 | 342 | 0.93 | 93 |
| problem/complication | 0.82 | 324 | 0.9 | 202 |
| future work | 0.83 | 869 | 0.85 | 195 |
| future prediction | 0.88 | 178 | 0.75 | 55 |
| important consideration | >0.99 | 735 | >0.99 | 491 |
| ALL CATEGORIES COMBINED* | 0.22 | 3354 | 0.12 | 2005 |

**Figure B.17:**
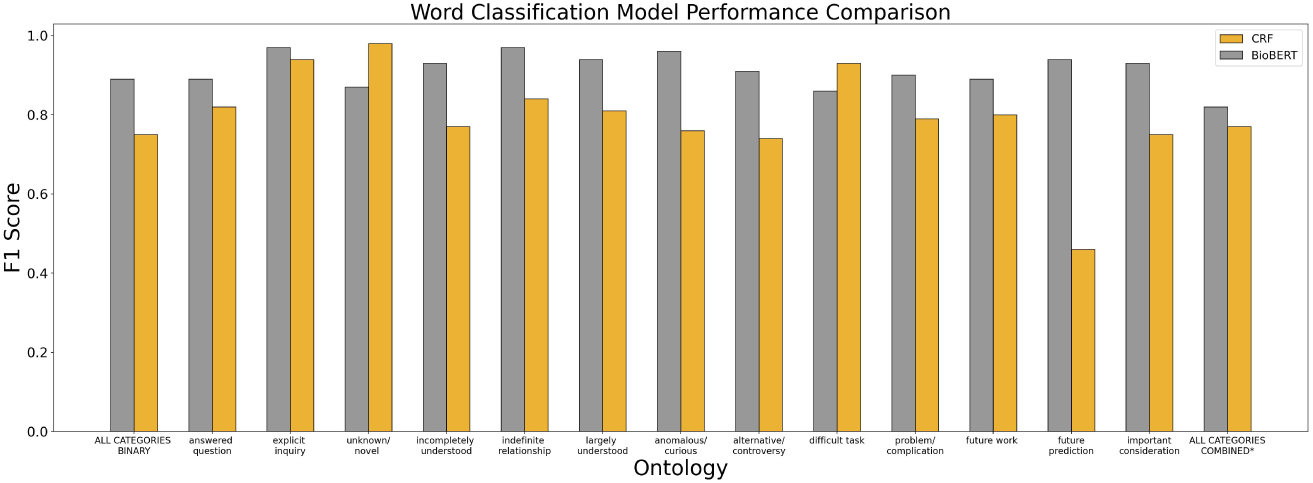
Word Classification summary: A bar plot summary of test F1 scores for word classification. *Reporting the macro-average F1 score of all the categories for one multi-classifier.

**Table B.17:** CRF word classification. *Reporting the macro-average F1 score of all the categories for one multi-classifier.

| Ignorance Category | training F1 score | training support | testing F1 score | testing support |
| --- | --- | --- | --- | --- |
| ALL CATEGORIES BINARY | 0.76 | 16748 | 0.75 | 7601 |
| answered question | 0.8 | 692 | 0.82 | 320 |
| unknown/novel | 0.97 | 401 | 0.98 | 155 |
| explicit inquiry | 0.94 | 136 | 0.94 | 43 |
| incompletely understood | 0.76 | 5764 | 0.77 | 2809 |
| indefinite relationship | 0.83 | 2847 | 0.84 | 1205 |
| largely understood | 0.79 | 1313 | 0.81 | 618 |
| anomalous/curious | 0.73 | 809 | 0.76 | 399 |
| alternative/controversy | 0.73 | 1262 | 0.74 | 598 |
| difficult task | 0.9 | 283 | 0.93 | 128 |
| problem/complication | 0.8 | 432 | 0.79 | 238 |
| future work | 0.82 | 991 | 0.8 | 391 |
| future prediction | 0.56 | 341 | 0.46 | 100 |
| important consideration | 0.75 | 1484 | 0.75 | 608 |
| ALL CATEGORIES COMBINED* | 0.77 | 13828 | 0.77 | 6239 |

**Table B.18:** BioBERT word classification. *Reporting the macro-average F1 score of all the categories for one multi-classifier.

| Ignorance Category | training F1 score | training support | testing F1 score | testing support |
| --- | --- | --- | --- | --- |
| ALL CATEGORIES BINARY | 0.9 | 16748 | 0.89 | 7601 |
| answered question | 0.92 | 692 | 0.89 | 320 |
| unknown/novel | 0.91 | 401 | 0.87 | 155 |
| explicit inquiry | 0.95 | 136 | 0.97 | 43 |
| incompletely understood | 0.94 | 5764 | 0.93 | 2809 |
| indefinite relationship | 0.98 | 2847 | 0.97 | 1205 |
| largely understood | 0.94 | 1313 | 0.94 | 618 |
| anomalous/curious | 0.91 | 809 | 0.96 | 399 |
| alternative/controversy | 0.9 | 1262 | 0.91 | 598 |
| difficult task | 0.9 | 283 | 0.86 | 128 |
| problem/complication | 0.88 | 432 | 0.9 | 238 |
| future work | 0.91 | 991 | 0.89 | 391 |
| future prediction | 0.97 | 341 | 0.94 | 100 |
| important consideration | 0.96 | 1484 | 0.93 | 608 |
| ALL CATEGORIES COMBINED* | 0.82 | 13828 | 0.82 | 6239 |

